# A novel recombinant PHB production platform in filamentous cyanobacteria avoiding nitrogen starvation while preserving cell viability

**DOI:** 10.1101/2024.10.01.616055

**Authors:** Phillipp Fink, Claudia Menzel, Jong-Hee Kwon, Karl Forchhammer

**Affiliations:** Organismic Interactions Department, Tübingen University, Auf der Morgenstelle 28, 72076 Tübingen, Germany; Division of Applied Life Sciences (BK21), Gyeongsang National University, Jinju 52828, Republic of Korea; Department of Food Science & Technology and Institute of Agriculture & Life Science, Gyeongsang National University, Jinju 52828, Republic of Korea

**Keywords:** Cyanobacteria, *Nostoc* sp. PCC7120, sustainable PHB production, Genetic engineering

## Abstract

During the past decades, the importance of developing sustainable, carbon dioxide (CO_2_)-neutral and biodegradable alternatives to conventional plastic has become evident in the context of global pollution issues. Therefore, heterotrophic bacteria such as *Cupriavidus* sp. have been intensively explored for the synthesis of the biodegradable polymer polyhydroxybutyrate (PHB). PHB is also naturally produced by a variety of phototrophic cyanobacteria, which only need sunlight and CO_2,_ thereby allowing a CO_2_ negative, eco-friendly synthesis of this polymer. However, a major drawback of the use of cyanobacteria is the need of a two-stage production process, since relevant amount of PHB synthesis only occurs after transferring the cultures to conditions of nitrogen starvation, which hinders continuous, large-scale production.

This study aimed at generating, by means of genetic engineering, a cyanobacterium that continuously produces PHB in large amounts. We choose a genetically amenable filamentous cyanobacterium of the genus *Nostoc* sp., which is a diazotrophic cyanobacterium, capable of atmospheric nitrogen (N_2_) fixation but naturally does not produce PHB. We transformed this *Nostoc* strain with various constructs containing the PHB synthesis operon from *Cupriavidus necator* H16. In fact, while the transformants initially produced PHB, the PHB-producing strains rapidly lost cell viability. Therefore, we next attempted further optimization of the biosynthetic gene cluster. Finally, we succeeded in stabilized PHB production, whilst simultaneously avoiding decreasing cell viability. In conclusion, the recombinant *Nostoc* strain constructed in the present work constitutes the first example of a continuous and stable PHB production platform in cyanobacteria, which has been decoupled from nitrogen starvation and, hence, harbours great potential for sustainable, industrial PHB production.

## Introduction

The production of non-degradable plastics from fossil oil poses a significant threat to global ecosystems (1,2). To ensure a sustainable future, it is crucial to prevent overproduction of conventional plastics from spiralling out of control. Instead, we must pursue alternative, bio- based and CO2 neutral pathways for the production of biodegradable polymers (3). Polyhydroxyalkanoates (PHA) are promising candidates to substitute petroleum-based plastics (4). PHA can be naturally produced and polymerized by various microorganisms (5,6). One of the most well-studied model organisms for bioplastic production is the heterotrophic bacterium *Cupriavidus necator* H16 (7).

*Cupriavidus necator* H16 (8), formerly known as *Hydromonas* sp. H16, *Alcaligenes eutrophus* H16 (9), *Ralstonia eutropha* H16 (10) and *Wautersia eutropha* H16 (11), has the ability to produce a variety of PHAs (7,12). The most widely studied PHA that has gained attention of scientist worldwide is polyhydroxybutyrate (PHB). In heterotrophic bacteria, PHB serves as an excessive carbon storage and is produced in large quantities under nutrient limited conditions (13). PHB can constitute up to 90 % PHB of the cell dry weight (CDW) (14,15). The biosynthesis of PHB in *Cupriavidus necator* H16 involves a three-enzyme pathway (Fig. 1).

**Fig. 1:**
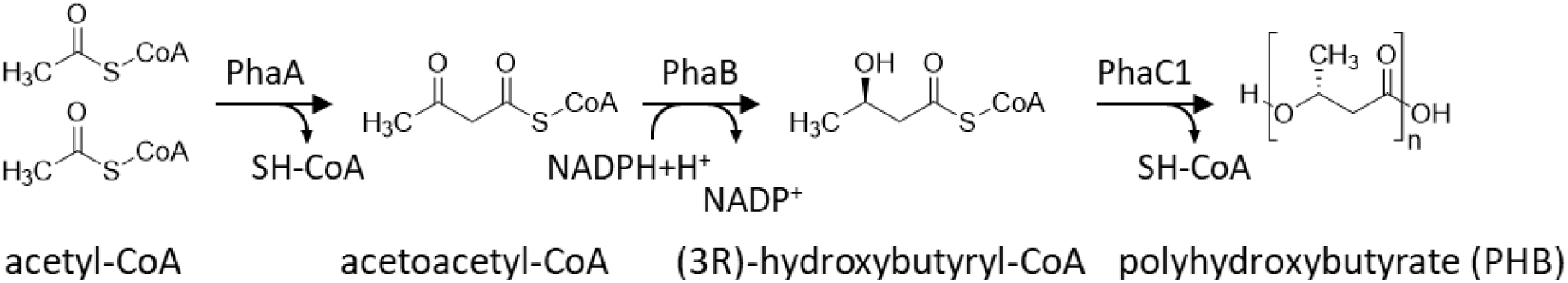
Biosynthesis of polyhydroxybutyrate (PHB) in *Cupriavidus necator* H16, two acetyl-CoA units were condensed into acetoacetyl-CoA by β-ketothiolase PhaA, acetoacetyl-CoA was reduced by acetoacetyl-CoA reductase PhaB to form 3-hydroxybutyryl-CoA, 3-hydroxybutyryl-CoA was polymerized by PHB synthase PhaC.

First, two acetyl-CoA units are condensed to form acetoacetyl-CoA by the β-kethothiolase PhaA. In the next step, acetoacetyl-CoA is reduced by the NADPH-dependent acetoacetyl-CoA reductase PhaB to form 3-hydroxybutyryl-CoA. Subsequently, the 3-hydroxybutyryl-CoA monomers are polymerized to PHB by the PHB synthase PhaC (16,17). In *Cupriavidus necator* H16, different isoforms of PhaC were reported (18). Mainly PhaC1 is active and responsible for PHB production by forming a dimeric, active structure on the surface of PHB granules (19). PHB granules consist of a spherical PHB core, which is coated by a layer of various PHB granule associated proteins (PGAP’s) to form a well-organized and complex structure termed “carbonosome” (20). Phasins are a particularly important class of PGAP’s. They are small, amphiphilic proteins that mediate between the hydrophobic inside of the PHB granulum and the hydrophilic, cytoplasmic periphery. Phasin PhaP1 is one of the most abundant PGAP’s in the carbonosome of *Cupriavidus necator* H16 (21). The absence of PhaP1 affects the overall PHB accumulation, resulting in a specific PHB mutant phenotype (22). PhaP is regulated by PhaR, which is also found bound to the PHB granule (23). Another phasin, PhaM, is responsible for the association and further activation of PhaC1, as well as for the correct distribution of PHB granules during cell division by binding to the chromosome (24,25). Other classes of PGAP’s that should be mentioned are PHB depolymerases, which are responsible for the PHB degradation (26). Investigating the structural assembly of the carbonosome is essential for understanding the complex regulatory machinery behind the PHB accumulation, which is crucial for industrial upscaling.

Currently, heterotrophic bacteria like *Cupriavidus necator* H16 and recombinant *E. coli* strain accumulating PHB are used for industrial, commercial PHB production (27–29). For heterotrophic PHB production, the bacteria need to be fed with organic carbon sources coupled with limited nutrient cultivation for efficient PHB production (30,31). However, the use of organic carbon sources, which alternatively might be used for nutrition, is highly expensive and not sustainable (32,33). Therefore, alternative processes have to be considered, such as bio-waste products instead of feeding high-value carbon sources (34). Alternatively, photoautotrophic synthesis of PHB using appropriate microorganism like cyanobacteria came into focus of research (35–39). Oxygenic photosynthesis by cyanobacteria utilizes sunlight and water for CO_2_ fixation, and thereby contributes to the decrease of the greenhouse gas CO_2_ in the atmosphere (40). Moreover, it avoids the consumption of organic carbon sources for PHB synthesis, if wastewater is used for the cultivation of cyanobacteria, an additional benefit for the environment can be achieved (41). This eco-friendly approach for PHB production is highly promising for a sustainable, PHB production.

PHB production by cyanobacteria was recorded since 1966, with the first report demonstrating the presence of PHB in *Chlorogloea fritschii* (42). To date, a plethora of several cyanobacteria were characterized producing PHB under phototrophic growth condition. However, only small percentages of PHB accumulation (< 3.5 % *(w/w)* PHB/ CDW) could be reported (43). Major PHB production rates were reported when cyanobacteria were transferred to medium with limited nutrient supply like nitrogen or phosphor deprivation (44).

Under nitrogen depleted condition, the unicellular cyanobacterium *Synechocystis* sp. 6803 achieved 9.5 % *(w/w)* PHB/CDW after 14 days were achieved. Additional phosphate limitation could increase PHB accumulation further to 11.2 % PHB/CDW (45). For genetically engineered *Synechocystis* sp. 6803, PHB production rates up to 63 % *(w/w)* PHB/CDW in optimized cultivation conditions could be reported (39). In the filamentous cyanobacterium *Nostoc muscorum*, PHB rates up to 8,6 % *(w/w)* were achieved under combined nitrogen and phosphorus depletion (46). To date, industrial upscale strategies for PHB production in cyanobacteria revolve around a two-stage cultivation system, where first biomass is accumulated in a vegetative growth phase with only low PHB synthesis, followed by the PHB accumulation phase under nutrient limited, growth arrested conditions (47). Two-stage cultivations have the major drawback of difficult implementation for upscale processes. Continuous PHB accumulation in cyanobacteria would be preferable for industrial upscale processes (48). However, during continuous, autotrophic growth condition, PHB synthesis in cyanobacteria did not reach the relevant of PHB yield to be competitive enough in comparison to heterotrophic PHB production.

This study aimed to overcome the problem of low PHB yield during autotrophic growth conditions of cyanobacteria by implementing the continuous PHB synthesis machinery from *Cupriavidus necator* into a genetically tractable cyanobacterium. The filamentous cyanobacterium strain *Nostoc* sp. PCC7120, formerly known as *Anabaena* sp. PCC7120 (49–51), was chosen due to the availability of molecular genetic tools developed during the past decades (52–55). In addition, the autoflocculation ability of *Nostoc* sp. PCC7120 holds great potential for cost-effective downstream harvesting in industrial applications (56). As *Nostoc* sp. PCC7120 cannot produce PHB naturally, the consequences of PHB overproduction can be specifically investigated due to the lack of regulatory mechanism of PHB biosynthesis common in natural PHB producer (53). Here, we report the step-by-step development of a recombinant *Nostoc* sp. PCC7120 strain that constantly produces PHB, paving the way for utilizing these organisms as a chassis for heterologous PHB production.

## Material & Methods

### Bacterial strains and growth conditions

*Escherichia coli* (*E. coli*) strains were grown in lysogeny broth (LB)(57), formulated as described by Lennox (5 g/L NaCl) at 37 °C, supplemented with appropriate antibiotics. For growth on solid media, 1.5 % (*w/v*) agar was added.

*Nostoc* sp. PCC7120, formerly known as *Anabaena* sp. PCC7120, and derived strains were grown in modified BG11 medium, supplemented with 5 mM NaHCO_3_ (50). More specifically, the formulation of BG11 was altered as follows: The amount of Na_2_CO_3_ was increased to 0.04 g/L, while ferric ammonium citrate was replaced by Fe(III)-citrate. If necessary, appropriate antibiotics were introduced to the culture. For nitrogen starvation, cells were grown in BG11 medium where NaNO_3_ was replaced by NaCl (BG11_0_). The cells were cultivated at 120 rpm and constant illumination of 30 - 40 µmol m^-2^s^-1^ at 28 °C. For cultivation on solid media, 1.5 % (*w/v*) Difco Agar was added. A list of the strains described in the present work is provided in Table S 1.

### Construction of mutant strains

Plasmids were constructed via Gibson assembly(58). DNA fragments were amplified with the Q5 High-Fidelity DNA Polymerase (NEB) and the primer listed in Table S 2. Isolation and purification of plasmid DNA and PCR product purification were conducted with the NEB Monarch Plasmid Miniprep Kit, the Monarch Gel Extraction Kit and the NEB Monarch PCR Purification Kit (New England Biolabs GmbH, Frankfurt am Main, Germany), according to manufacturer specifications. All constructs were verified by restriction digest and sequencing (Eurofins Genomics, Ebersberg, Germany) and are listed in Table S 3. Plasmids were then introduced into *Nostoc* via triparental conjugation (52). The subsequent screening for double-crossover mutants was performed by plating exconjugants on BG11 agar plates, containing the respective antibiotics and a 5% (*w/v*) sucrose concentration. Single crossover mutants integrated the whole plasmid containing the selection gene *sacB* are not able to grow in presence of sucrose.

### Fluorescence microscopy

Fluorescence microscopy was performed on a LeicaDM5500B (Leica Microsystems, Germany) with objective lenses and filter cubes listed in Table 1:

**Table 1:** LeicaDM5500B microscope components.

| Objective lenses | Magnification | Numerical aperture | Immersion |
| --- | --- | --- | --- |
| HCX PL FLUOTAR 40x/0.75 DRY | 40 x | 0.75 | Air |
| HCX PL FLUOTAR 100x/1.30 OIL | 100 x | 1.30 | Oil |
| Filter cubes | Excitation Filter | Dichromatic Mirror | Emission Filter |
| GFP | BP 470/40 | 500 | BP 525/50 |
| CY3-T | BP 560/10 | 560 | 610/65 |

BODIPY signal detection was performed using the GFP channel, while cyanobacterial autofluorescence was detected by using the CY3 channel.

### PHB granule staining procedure

PHB granules within cells were stained with the fluorescent dye BODIPY. For this purpose, 1 µL of a BODIPY solution (1 mg/ml dissolved in DMSO) was added to 100 µL of cyanobacterial culture. After 5 min of incubation in the dark, cells were centrifuged (3500 ×g, 5 min, RT), washed with 100 µL PBS buffer and resuspended in 10 µL PBS buffer. Then, the resuspended cells were applied on microscopic slides coated with a mixture of 1 % (*w/v*) agarose and 0.05 mg/ml propyl- cyanophycin for cell immobilization.

### Electron microscopy

Sample preparation was performed as described by Fiedler et al. 2002. Briefly, *Nostoc* sp. samples were fixed and post-fixed with 25 % (*w/v*) glutaraldehyde and 2 % (*w/v*) potassium permanganate, respectively. After embedding in EPON and staining with 2 % (*w/v*) uranyl acetate and lead citrate, samples were examined with a Philips Tecnai 10 electron microscopy at 80 kV.

### PHB quantification

The method described in (59,60) was optimised and the PHB content of cyanobacterial strains was determined as follows. 12 - 15 ml bacterial culture was harvested by centrifugation (4000 g, 10 min, RT) in pre-weighed reaction tubes and dried overnight at 90 °C. Cell dry weight was determined, and the cells were subsequently boiled in 1 ml of concentrated H_2_SO_4_ (18 M) for 1 h at 100 °C in order to convert PHB to crotonic acid. Then, 100 µL of boiled cell culture was diluted with 900 µL 0.014 M H_2_SO_4_ and centrifuged at 20,000 g for 5 min at RT. Next, 500 µL of the supernatant was mixed with 500 µL 0.014 M H_2_SO_4_ and centrifuged again. The resulting supernatant was analysed by high-pressure liquid chromatography (HPLC) (HITACHI Chrommaster, VWR, Germany). 5 µL of samples were injected onto a reverse phase column (Nucleosil 100-5 C18 column, particle size 5 µm, pore size 100 Å, 125 x 3 mm, fitted with a precolumn 4 x 3 mm) at a flowrate of 1 ml/min and eluted with an isocratic mixture of 30/70 MeOH/20 mM phosphate buffer (pH = 2.5) over 10 min for crotonic acid detection. Detection and crotonic acid quantification were conducted at λ = 210 nm. PHB samples with known concentrations were prepared and processed as described above to determine the conversion rate of PHB to crotonic acid. Commercially available crotonic acid was measured in defined concentrations for linear regression curve fitting.

## Results

### Construction of first generation recombinant *Nostoc* sp. PCC7120 strain NosPHB1.0

To prove the suitability of *Nostoc* sp. PCC7120 as a PHB production host, the PHB synthesis operon from *Cupriavidus necator* H16 containing the genes *phaCAB* was placed under the control of the constitutive promoter P*_pbsA_* from *Amaranthus hybridus* (61) and cloned into the replicative plasmid pRL1049. Then, the resulting plasmid pRL1049-P*_psbA_*-*phaCAB* was introduced into the filamentous organism by triparental mating., using the helper strain *E. coli* J53/RP4 and the cargo strain *E. coli* HB101/pRL528/pRL1049-P_psbA_-*phaCAB,* resulting in the recombinant strain NosPHB1.0. After incubation on agar plates, the obtained exconjugants were examined by fluorescence microscopy to visualize PHB granules using BODIPY green straining and determine the distribution of stained PHB granules. (Fig. 2)

**Fig. 2:**
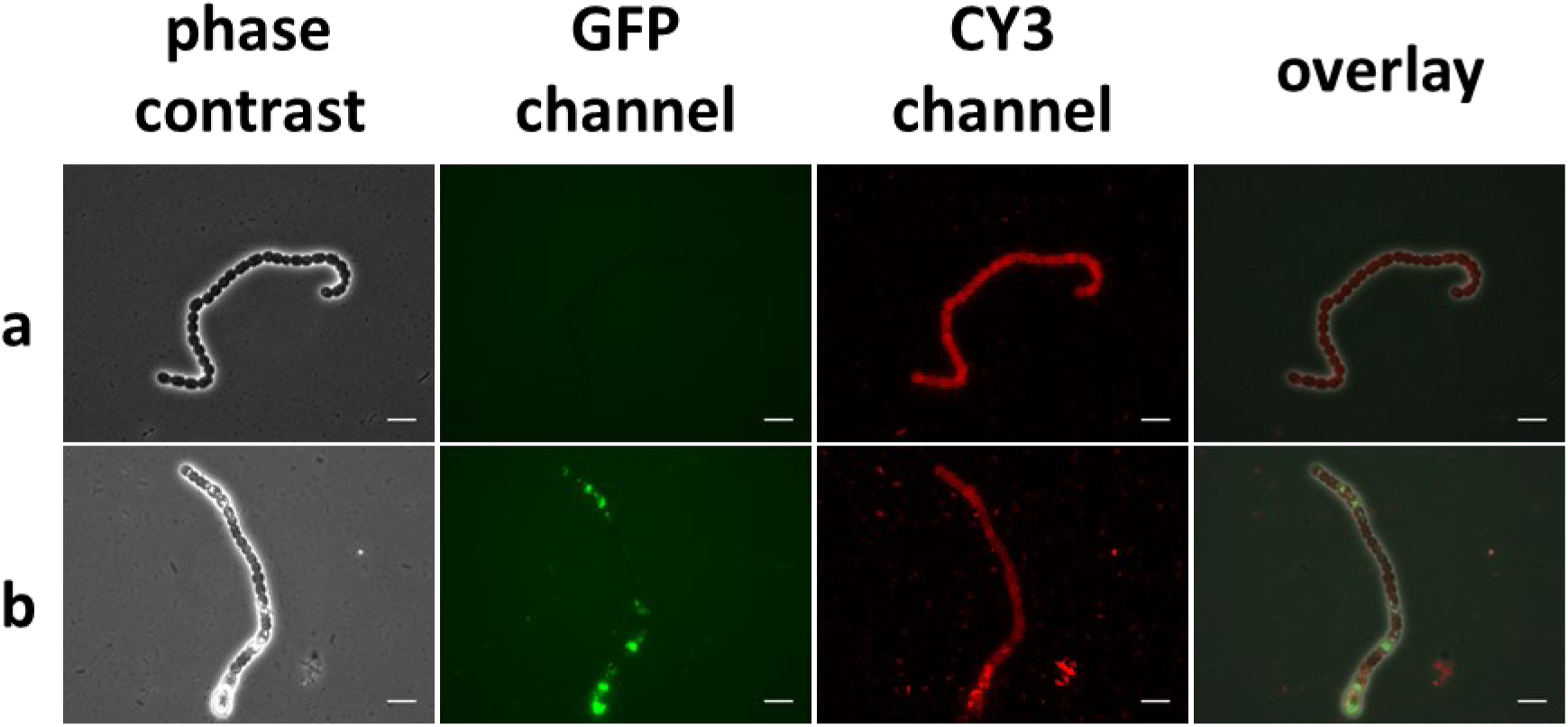
Microscopic images of wild type and NosPHB1.0 strain **(a)** Wildtype *Nostoc* sp. 7120 **(b)** Recombinant *Nostoc* strain NosPHB1.0. PHB granules are visualized with BODIPY staining (GFP channel), autofluorescence (CY3 channel), overlay of phase contrast, GFP and CY3 channel, scale bar = 10 µm.

Indeed, PHB granules could be detected inside some cells of filaments, as shown exemplary in the microscopic image in Fig. 2.. However, PHB granules were not equally distributed in the individual cells but happened to arise heterogeneously in the observed filament, with some cells displaying strong BODIPY straining, while other appeared empty. A similar phenotype of potential PHB granule production could be observed in all exconjugants by fluorescence microscopic examination (data not shown).

### Initial characterization of PHB accumulation in NosPHB1.0

To validate that the observed BODIPY-stained granules consist indeed of PHB and to gain first insights into the recombinant PHB producer strain NosPHB1.0, initial cultivation experiments and first PHB quantifications were performed with one recombinant strain as a proof of principle (Fig. 3).

**Fig. 3:**
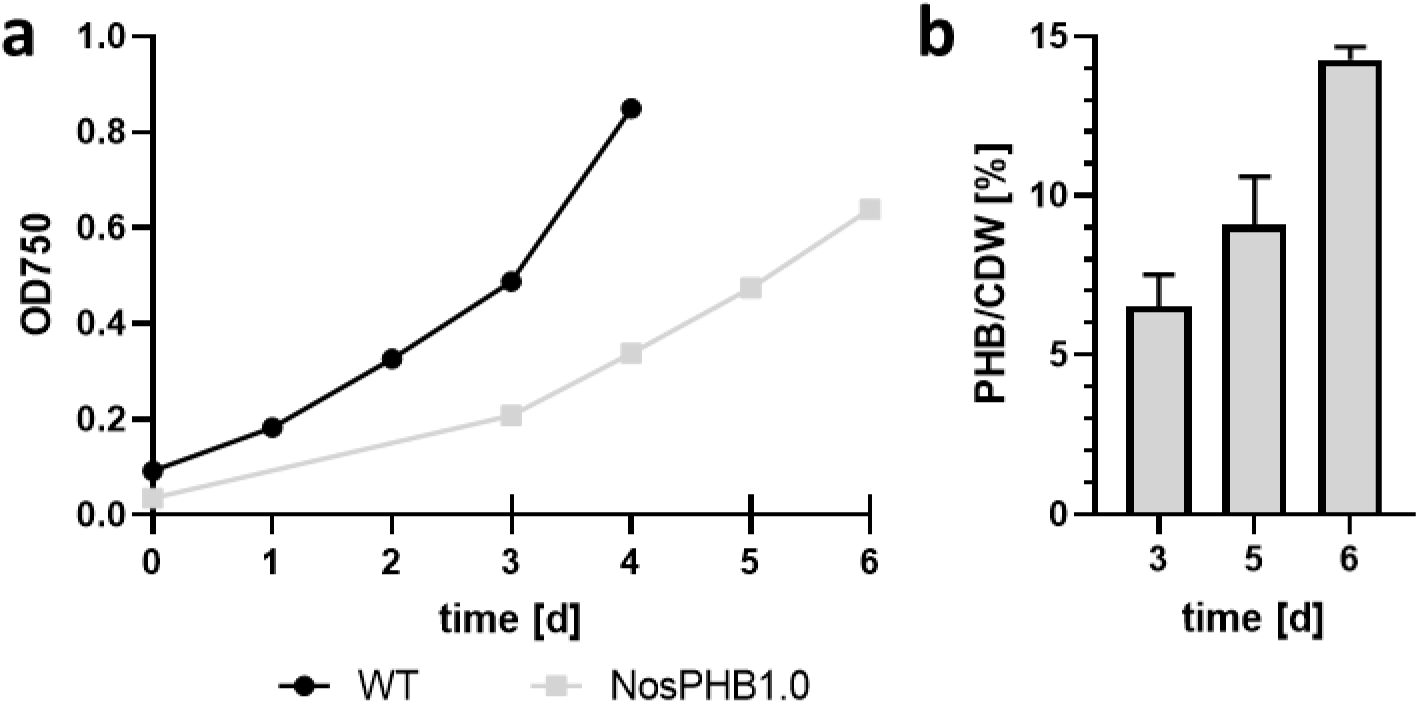
Characterization of NosPHB1.0 **(a)** Growth curve of wild-type strain *Nostoc* sp. PCC7120 (black) and NosPHB1.0 (gray), recorded by measuring the OD_750_. Longer ticks on the x-axis indicate samples taken for PHB quantification. Each point represents one biological replicate **(b)** PHB content of NosPHB1.0 after 3, 5 and 6 days of constant growth conditions. Each data set represents duplicates of the same biological sample.

Under standard growth conditions, NosPHB1.0 showed a slower growth in comparison to the wild-type *Nostoc* sp. PCC7120 (Fig. 1, a). Initial PHB content measurement showed that up to 14 % *(w/w)* PHB/CDW was achieved after six days of phototrophic cultivation in BG11 medium (Fig. 1, b). These data confirmed that NosPHB1.0 is the first recombinant PHB producing Nostoc strain.

### Morphological change of NosPHB1.0

In order to examine PHB accumulation in the recombinant *Nostoc* strain and correlate microscopic images with HPLC quantification, samples for fluorescence microscopy were taken in parallel to samples for PHB quantification. Samples from microscopic analysis were stained with BODIPY (Fig. 4) and microscopic images were taken in the GFP channel to visualize the green fluorescence coming form PHB-embedded BODIPY and in the CY3 channel to visualize autofluorescence from the photosynthetic pigments of vegetative cells.

**Fig. 4:**
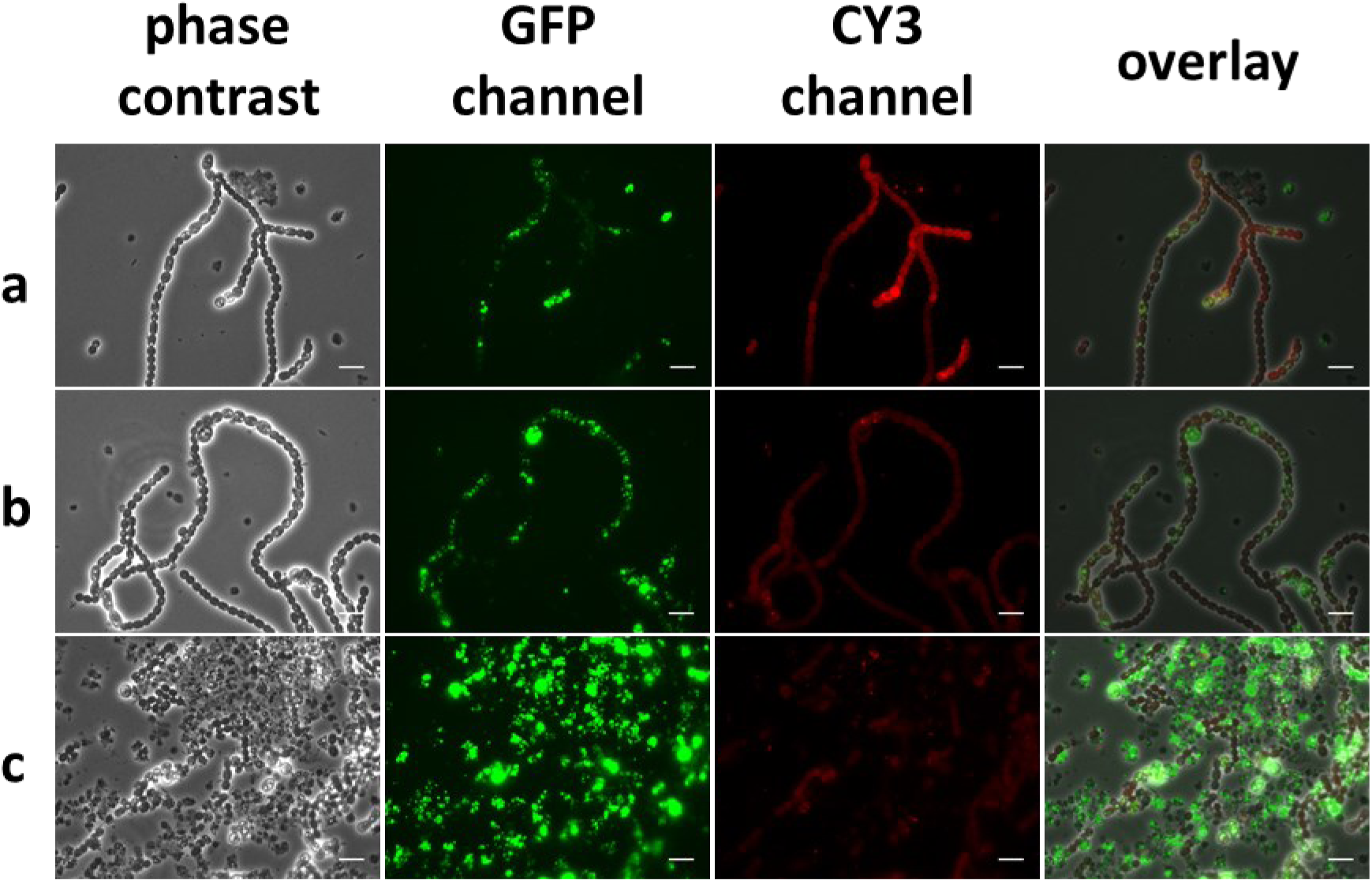
Microscopic images of NosPHB1.0 after **(a)** 3 days, **(b)** 5 days and **(c)** 6 days of constant cultivation conditions, PHB granules are visualized with BODIPY staining (GFP channel), autofluorescence (CY3 channel), overlay of phase contrast, GFP and CY3 channel, scale bar = 10 µm.

Strikingly, PHB accumulation occurred heterogeneously in the filament during the entire course of the experiment, while remarkable growth morphologies of NosPHB1.0 could be noted (Fig. 4). At the start of the cultivation, recombinant filaments began to grow and accumulate PHB heterogeneously (Fig. 4, a). PHB accumulation increased over time in the filaments (Fig. 4 b). However, during prolonged cultivation, cells accumulating PHB got increasingly detached from main filaments, resulting in single cells displaying the highest PHB content. Furthermore, these cells showed reduced autofluorescent signals, indicating loss of photosynthetic capacity (Fig. 4, c). We termed these accumulations of PHB-filled single cells “PHB graveyards”.

### NosPHB1.0 grown under different growth conditions

For further characterization of the recombinant *Nostoc* strain, preliminary experiments under different shaking conditions and light intensities were tested in single cultivation experiments to determine optimal PHB production conditions for NosPHB1.0 (Fig. 5).

**Fig. 5:**
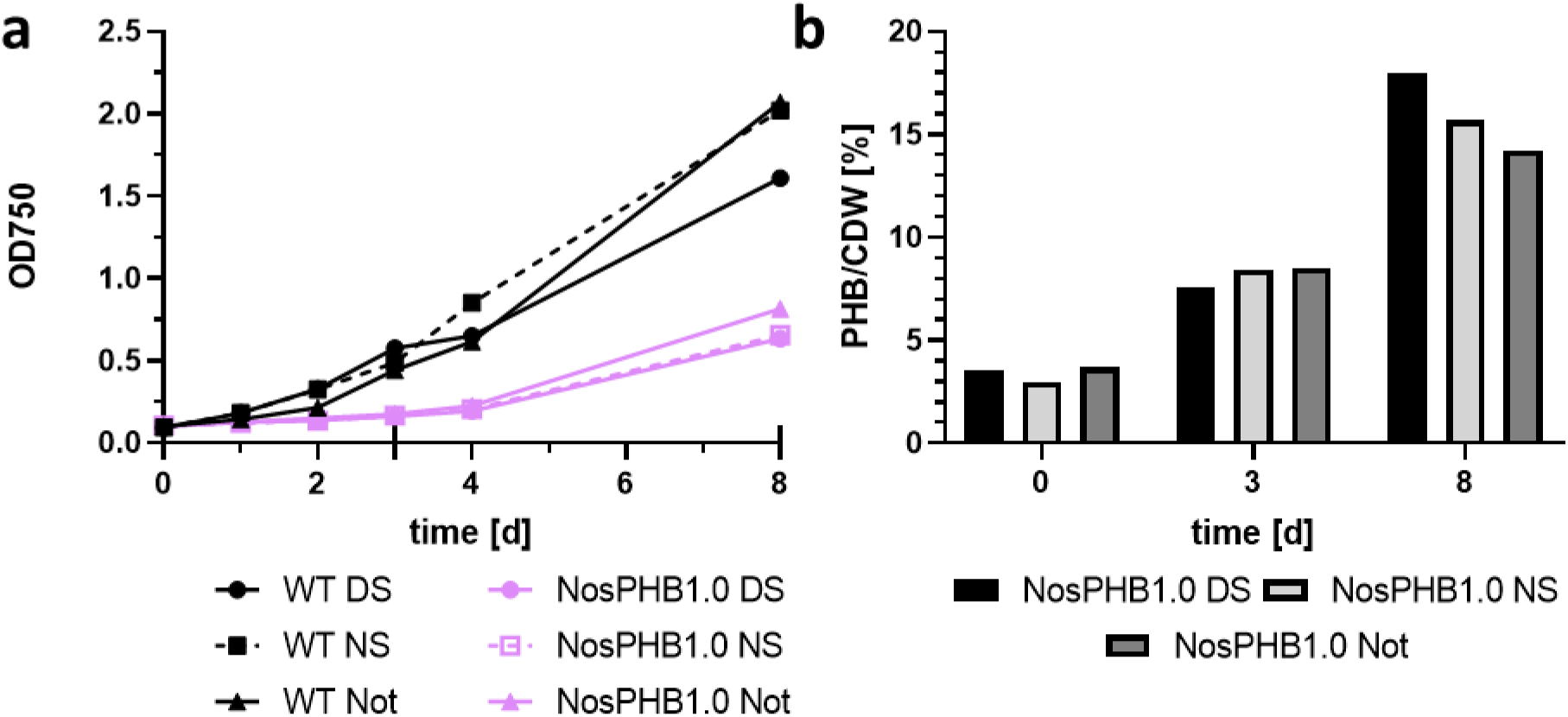
Growth curve and PHB content of NosPHB1.0 measured under different conditions. **(a)** Growth curve of wild-type strain *Nostoc* sp. PCC7120 (black) and NosPHB1.0 (violet) under various growth conditions, DS: 20 µmol m^-2^s^-1^, 120 rpm; NS: 40 µmol m^-2^s^-1^, 120 rpm; Not: 20 µmol m^-2^s^-1^, 120 rpm, recorded by measuring the OD_750_. Longer ticks on the x-axis indicate samples taken for PHB quantification. Each point represents one biological replica recorded by measuring the OD_750_. Longer ticks on the x-axis indicate samples taken for PHB quantification. Each point represents one biological replicate**(b)** PHB content of NosPHB1.0 after 0, 3 and 8 days of cultivation under various growth conditions. Each data set represents one biological sample.

Cultures were either cultivated under dimmed light of 20 µmol m^-2^s^-1^ and constant shaking at 120 rpm (DS), constant light illumination of 40 µmol m^-2^s^-1^ and shaking at 120 rpm (NS) or dimmed light condition of 20 µmol m^-2^s^-1^ and no shaking (Not) (Fig. 5, a), measuring the optical density (OD_750_) as a proxy for cell-growth and quantification of PHB content. Compared to the wild type strain, NosPHB1.0 displayed drastically reduced growth rates, supporting our previous finding regarding NosPHB1.0 in biological triplicates. NosPHB1.0, when cultivated without shaking showed a slightly higher growth rate compared to the other conditions (Fig. 5, a). All three conditions showed similar PHB/CDW rate at the beginning of the experiment. However, the highest PHB/CDW rate was obtained after 8 days of dimmed light cultivation “DS”, where up to 17 % *(w/w)* PHB/CDW was measured (Fig. 5, b). However, prolonged cultivation of NosPHB1.0 led to a separation of phenotypes, consisting of non-PHB producing filaments and PHB containing single cells (Fig. 6).

**Fig. 6:**
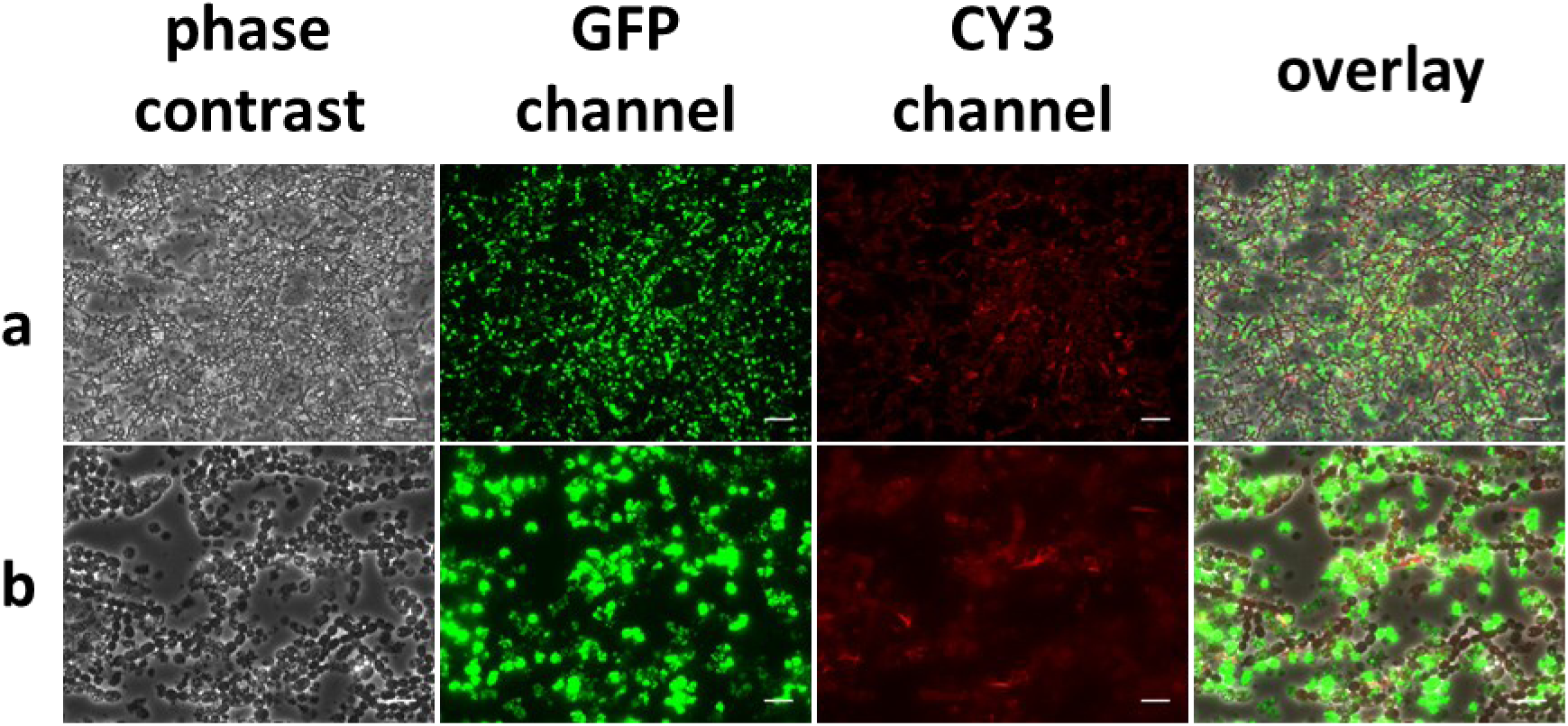
Microscopic images of NosPHB1.0 after 18 days of continuous cultivation, PHB granules are visualized with BODIPY staining (GFP channel), autofluorescence (CY3 channel), overlay of phase contrast, GFP and CY3 channel, **(a)** scale bar = 25 µm, **(b)** scale bar = 10 µm.

These experiments reinforced the previous observations. While the number of PHB producing filaments decreased, most PHB granula were detected in apparently deteriorating single cells as seen by the accumulation of “PHB graveyards” (Fig. 6). A potential reason explaining the loss of PHB synthesis in entire filaments could be the partial loss of the replicative plasmid encoding for the PHB operon. Similar cases of plasmid expulsion in cyanobacteria have been noticed in former studies (Castets et al. 1986). Therefore, to avoid the potential loss of the replicative plasmid and the subsequent lack of PHB production inside the filaments, a novel cloning strategy was devised by integrating the PHB operon in the genome of *Nostoc* sp. PCC7120.

### Second generation recombinant *Nostoc* sp. PCC 7120 PHB producer NosPHB2.0

In order to integrate the PHB operon into the genome of *Nostoc* sp. PCC7120, an integrative plasmid for triparental mating and subsequent recombination into the known *nucA-nuiA* neutral site of *Nostoc* (62) genome was constructed. For this purpose, plasmid pPF08 was constructed: The backbone of pRL271 was combined with sequences homologous to the *nucA-nuiA* neutral site flanking the modified PHB operon, which consists of the constitutive promotor P_psbA_, the biosynthetic genes *phaCAB* from *Cupriavidus necator* H16 and the selection marker, *aad1*, which encodes for an adenylytransferase, enabling the selection of successful recombination events with spectinomycin and streptomycin. To prevent downstream effects of the constitutive promoter, a t_1_t_2_ terminator was added at the 3’end of the operon. After verification by sequencing, pPF08 was introduced into *Nostoc* PCC 7120 via triparental mating with helper strain *E. coli* J53/RP4 and cargo strain *E. coli* HB101/pRL528/pPF08. The successful double-crossover event and thus integration into the genome was validated by colony PCR and sequencing. Microscopic pictures were taken before and after screening for double-crossover mutants. (Fig. 7)

**Fig. 7:**
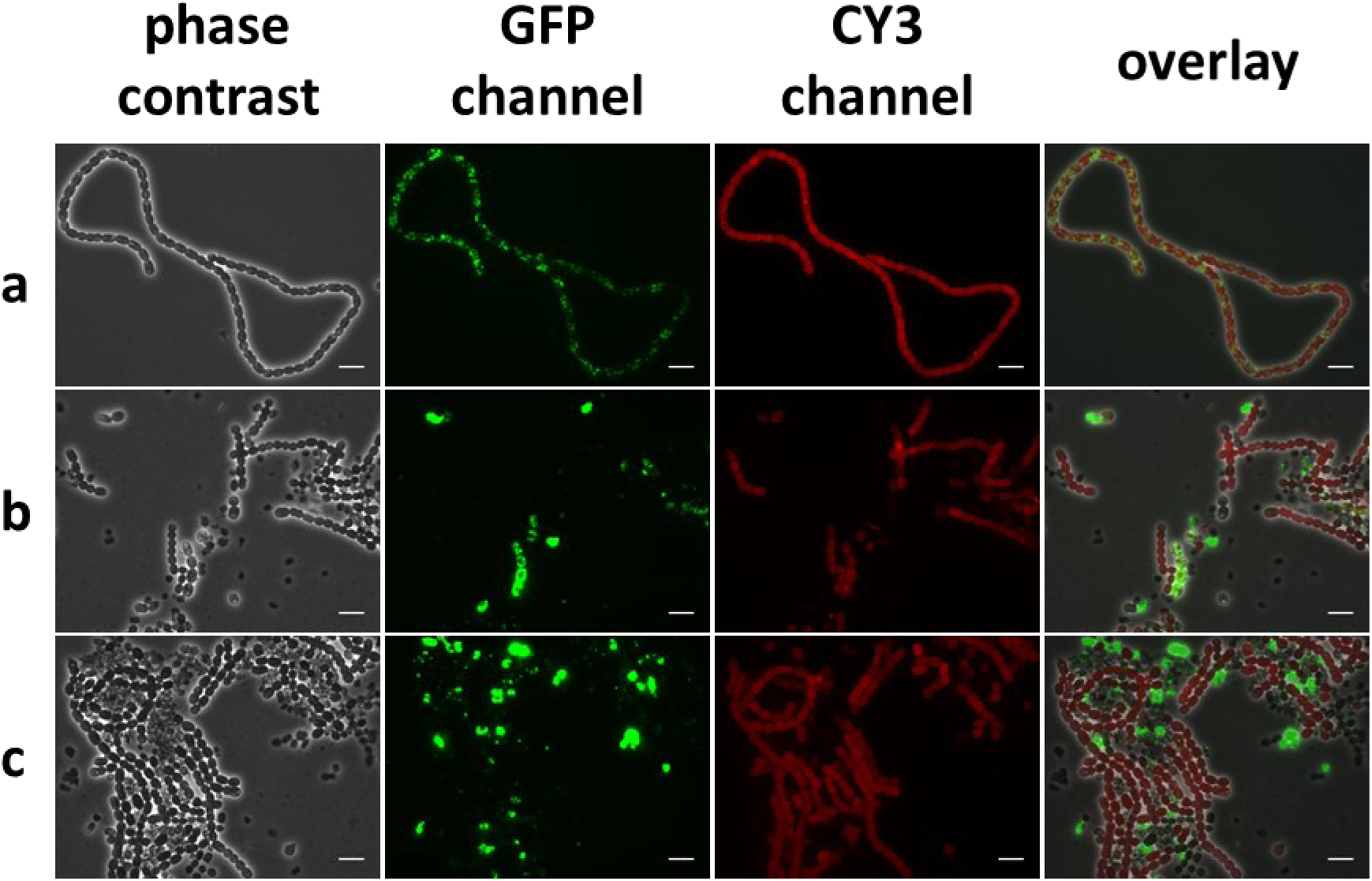
Microscopic images of NosPHB2.0 of single colonies **(a)** before double crossover screening, **(b)** after double crossover screening and **(c)** after 8 days of continuous cultivation. PHB granules are visualized with BODIPY staining (GFP channel), autofluorescence (CY3 channel), overlay of phase contrast, GFP and CY3 channel, scale bar = 10 µm.

Prolonged screening for double-crossover mutants via sucrose selection resulted in a delay of approximately 21 days before first cultivation experiments could be performed, in contrast with the more rapid screening for positive mutants in conjugation experiments with replicative plasmids. Exemplary microscopic pictures from exconjugants showed a more distributive, uniformly PHB production in NosPHB2.0 (Fig. 7, a) in comparison of NosPHB1.0. After double crossover screening of positive clone, PHB production could no longer be detected in all recombinant filaments (Fig. 7, b) while single cells, detached from the filament and filled with PHB like in the previously observed PHB graveyards became apparent. While biomass was increasing, PHB accumulation almost vanished. Simultaneously, prominent PHB signals were only observed in the single cells of the reoccurring PHB graveyards (Fig. 7, c). We concluded that the prolonged screening time for double recombinant exconjugants and therefore longer cultivation time, led to the loss of PHB production in the recombinant *Nostoc* strain once again. This was further confirmed in test cultivations of NosPHB2.0 in BG11 and BG11_0_, where PHB values of 5 % *(w/w)* PHB/CDW were achieved after 7 days of cultivation, which further decreased after 21 days to only 0.5 % *(w/w)* PHB/CDW (Fig. S 1). Thus, the previously proposed loss of the replicative plasmid harbouring the genes for PHB biosynthesis fails to explain the loss of PHB production during cultivation. Instead, the accumulation of PHB might exert negative effects on cell viability, resulting in a strong selective pressure to eliminate PHB production. We hypothesized that the hydrophobic surface of the recombinantly produced PHB might interfere with cellular structures such as thylakoid membranes that harbour the photosynthetic machinery. By contrast, natural PHB synthesizing bacteria contain specific PHB-surface shielding proteins, the amphiphatic phasines of the PhaP class (21). To test this hypothesis, we attempted a third strategy, where heterologous *phaP* gene was co-expressed with the PHB biosynthesis operon.

### Third generation recombinant *Nostoc* sp. PCC 7120 PHB producer NosPHB3.0

To potentially prevent toxic effects of heterologojusly produced PHB biosynthesis, the PHB operon in pPF08 was expanded with phasin gene *phaP1* from *Cupriavidus necator* H16. PhaP1 was reported to constitute the major PHB coating phasin in *Cupriavidus necator* (21). To ensure efficient translation of the phaP1 transcript, the native ribosomal binding sequence of the highly expressed *apcB* and *apcA* (*apcBA*) genes (63) was cloned in front of the phaP1 translational start site. The new construct, pPF10, was introduced into *Nostoc* sp. PCC7120 by triparental mating as described above for NosPHB2.0. Successful integration into the genome was validated by colony PCR and sequencing. To compare cell viability of the strains, the triparental mating of pPF08 and the subsequent recreation of NosPHB2.0 was performed in parallel. In order to avoid prolonged screening times causing loss of PHB production, screening of double-crossover mutants was neglected this time.

### Characterization of PHB accumulation in NosPHB3.0

NosPHB2.0 and NosPHB3.0 were grown in a continuous “seed” culture, where growth behaviour and PHB accumulation were observed over a prolonged period of cultivation time (Fig. 8). A benefit of choosing this seed culture cultivation strategy constitutes the repeated testing of the strain in each culture since every time the medium is refilled, the strain needs to adapt. Therefore, the seed culture provides a means of internal validation.

**Fig. 8:**
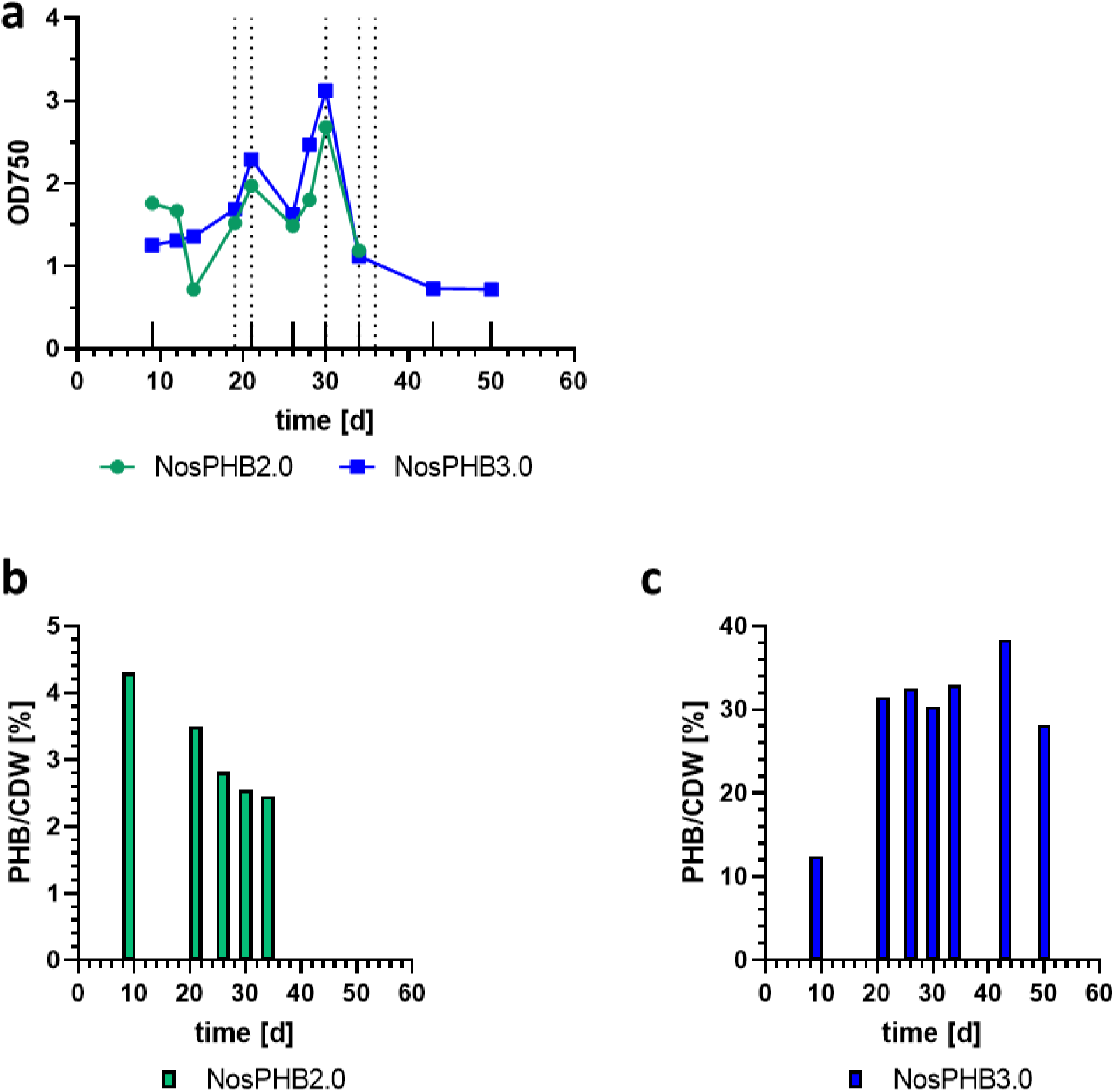
Growth experiment and PHB quantification of “seed” cultures of NosPHB2.0 and NosPHB3.0 **(a)** Growth curve of NosPHB2.0 (green) and NosPHB3.0 (blue), recorded by measuring the OD_750_. Longer ticks on the x-axis indicate samples taken for PHB quantification. Each point represents one biological replica recorded by measuring the OD_750_. Longer ticks on the x-axis indicate samples taken for PHB quantification. quantification, dashed lines indicate the refilling of BG11 medium to the original volume with appropriate antibiotics. Each point represents one biological experiment. **(b+c)** PHB content of NosPHB2.0 (green, **b**) and NosPHB3.0 (blue, **c**) after the respective days of continuous growth conditions. Each data set represents one biological sample.

For the following experiments, samples of the bacterial seed cultures were withdrawn, and the volume was refilled with BG11 medium with the respective antibiotics (Fig. 8, a). Over the course of this experiment, PHB/ CDW values above 30 % (*w*/*w*) were observed in the seed culture of NosPHB3.0 (Fig. 8, c). At the same time, PHB production rates in NosPHB2.0 decreased from an initial 4.3 % (*w*/*w*) PHB/CDW to 2.5 % (*w*/*w*) PHB/CDW in the span of 30 days (Fig. 8, b). In addition to the observed PHB content, microscopic pictures were taken from NosPHB2.0 and NosPHB3.0 (Fig. 9) corroborating the quantitative PHB analysis.

**Fig. 9:**
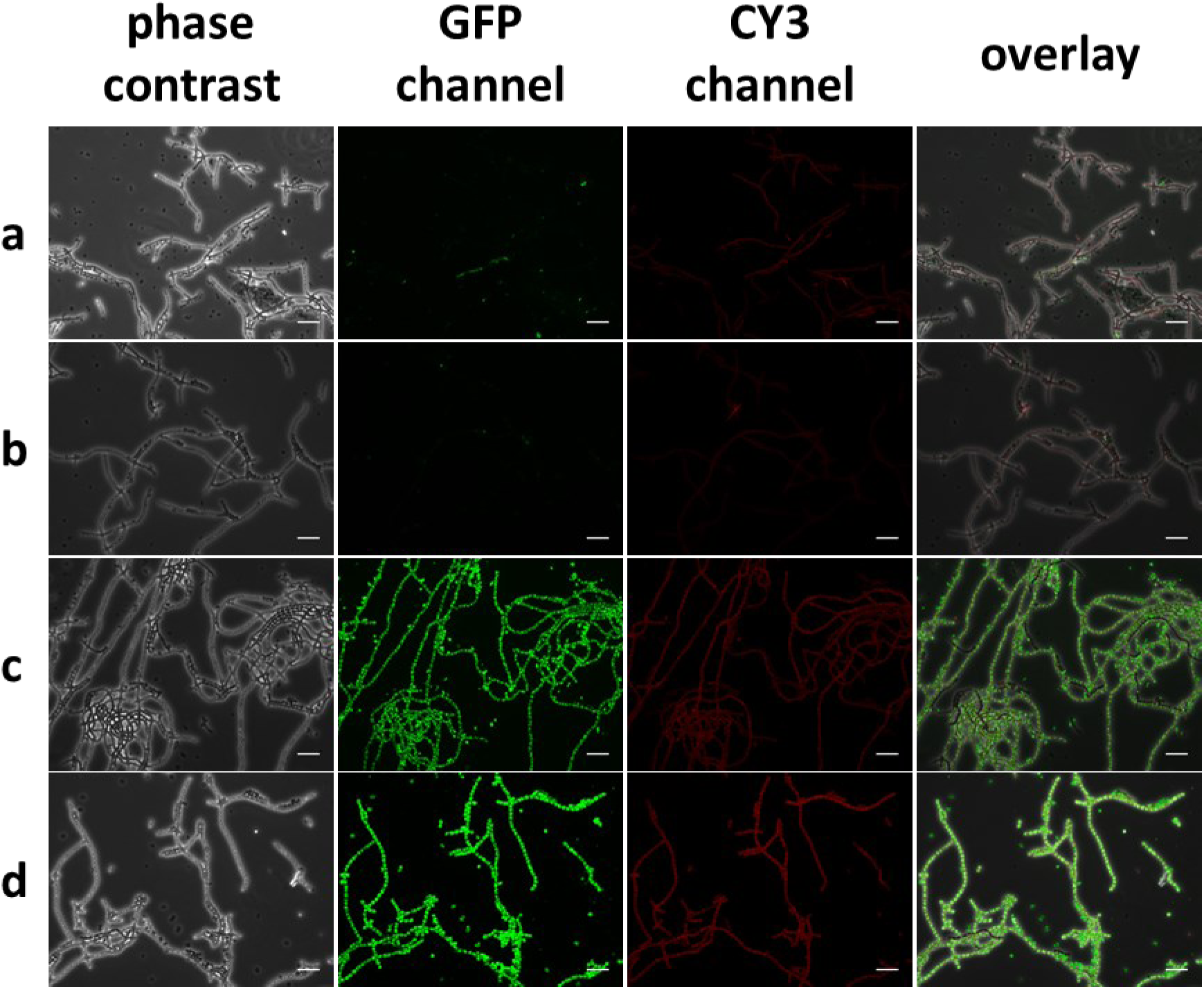
Microscopic images of **(a+b)** NosPHB2.0 and **(c+d)** NosPHB3.0 after **(a+c)** 12 days and **(b+d)** 30 days of continuous cultivation, PHB granules are visualized by BODIPY staining and detected using the GFP channel, PHB granules are visualized with BODIPY staining (GFP channel), autofluorescence (CY3 channel), overlay of phase contrast, GFP and CY3 channel, scale bar = 25 µm.

The signal for PHB granules in filaments of NosPHB2.0 was mostly abolished again during the prolonged cultivation (Fig. 9, a+b), supporting the previous findings. However, in NosPHB3.0, bright fluorescent PHB granules were detected, which were evenly distributed over the entire filaments. Moreover, the abundance of the fluorescent granules remained constantly hight during the continuous cultivation for 30 days (Fig. 9, c+d). This strongly implied that co-expression of phasin PhaP1 apparently stabilized PHB production over prolonged time periods.

In order to further characterize and subsequently optimize PHB production in NosPHB3.0, different shaking conditions and light intensities were tested. Cultures were either cultivated under dimmed light of about 5 µmol m^-2^s^-1^ and no shaking (NS), constant moderate illumination of 50 - 60 µmol m^-2^s^-1^ and shaking at 120 rpm (S) or dimmed light of 20 µmol m^-2^s^-1^ and constant shaking at 120 rpm (DS) (Fig. 10).

**Fig. 10:**
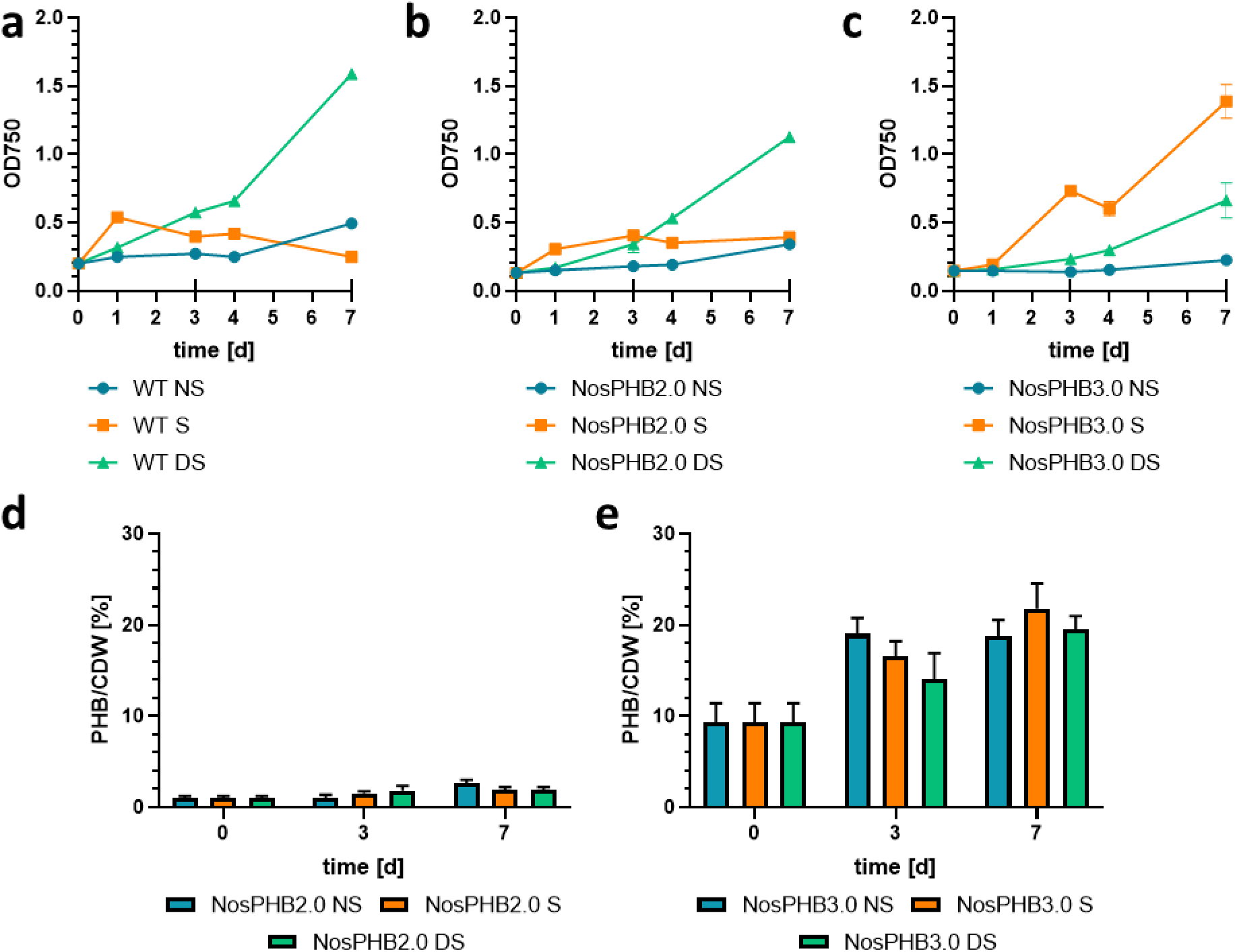
Growth curve of the wild-type strain, NosPHB2.0 and NosPHB3.0 and PHB production quantified in the latter two strains under different conditions. Growth curve of **(a)** Wild-type strain *Nostoc* sp. PCC7120, **(b)** NosPHB2.0 and **(c)** NosPHB3.0 under various growth conditions, NS: 5 µmol m^-2^s^-1^, 0 rpm; S: 50-60 µmol m^-2^s^-1^, 120 rpm; DS: 20 µmol m^-2^s^-1^, 120 rpm, recorded by measuring the OD_750_. Longer ticks on the x-axis indicate samples taken for PHB quantification. Each point represents one biological replica recorded by measuring the OD_750_. Longer ticks on the x-axis indicate samples taken for PHB quantification. Each data point represents **(a)** one, **(b)** two, **(c)** three biological replicates. **(d)** PHB content of NosPHB2.0 after 0, 3 and 7 days under various growth conditions. Each data set represents two biological replicates **(e)** PHB content of NosPHB3.0 after 0, 3 and 7 days of various growth conditions. Each data set represents three biological replicates.

While the growth curve of NosPHB2.0 resembled the wild-type strain (Fig. 10,a,b), NosPHB3.0 achieved the highest growth rate under constant illumination and shaking (condition S) (Fig. 10, c), as well as the highest PHB/CDW value of over 20 % (*w*/*w*) PHB/CDW after 7 days of cultivation (Fig. 10, e). In parallel, microscopic images of the cultivated NosPHB2.0 and NosPHB3.0 strains were taken (Fig. 11, Fig. S 2 Fig. S 5).

**Fig. 11:**
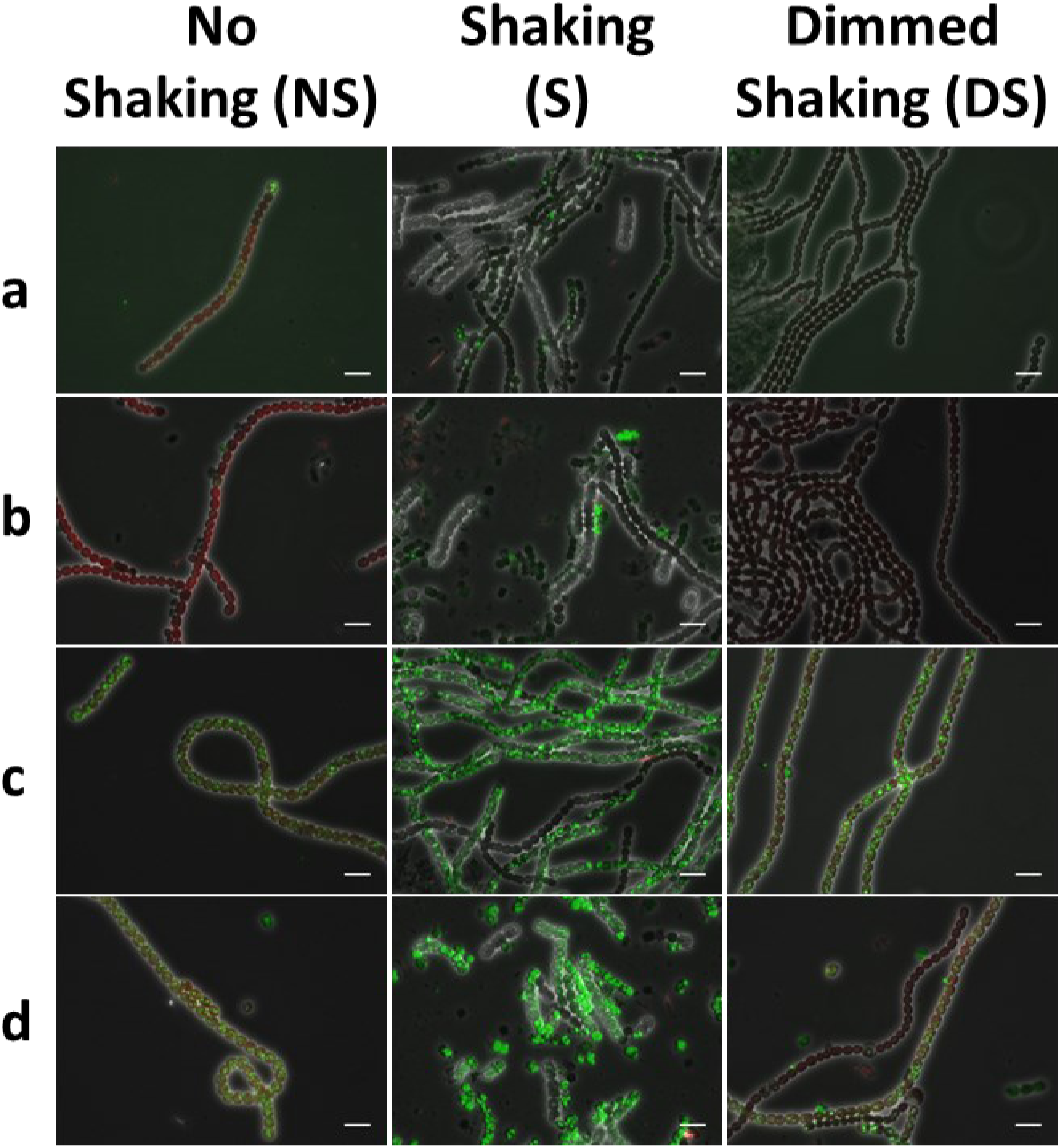
Overlay of microscopic images of **(a+b)** NosPHB2.0 and **(c+d)** NosPHB3.0 under various growth conditions after **(a+c)** 3 days and **(b+d)** 7 days of cultivation. PHB granules are visualized with BODIPY staining (GFP channel), autofluorescence (CY3 channel), overlay of phase contrast, GFP and CY3 channel, scale bar = 10 µm.

Similar as described above for one culture of NosPHB2.0, two independent cultures (NS and DS) of NosPHB2.0 displayed decreased PHB signals under all tested growth conditions (Fig. 11, a+b).

However, under condition S, small filaments with remaining PHB signals could be observed (Fig. 11, b). Filaments of NosPHB3.0 showed high fluorescence signals and, therefore, high accumulation of PHB (Fig. 11, c+d). Similar to NosPHB2.0, the highest PHB accumulation in NosPHB3.0 occurred under shaking condition with moderate light (Fig. 11, c+d).

### Effects of phasin PhaP1 on PHB granules visualised by transmission electron-microscopy (TEM)

In order to visualize the effects of phasin PhaP1 utilization in the recombinant *Nostoc* strain, TEM pictures were taken of *Nostoc* sp. PCC7120, NosPHB2.0 and NosPHB3.0 after 12 days of cultivation (Fig. 12).

**Fig. 12:**
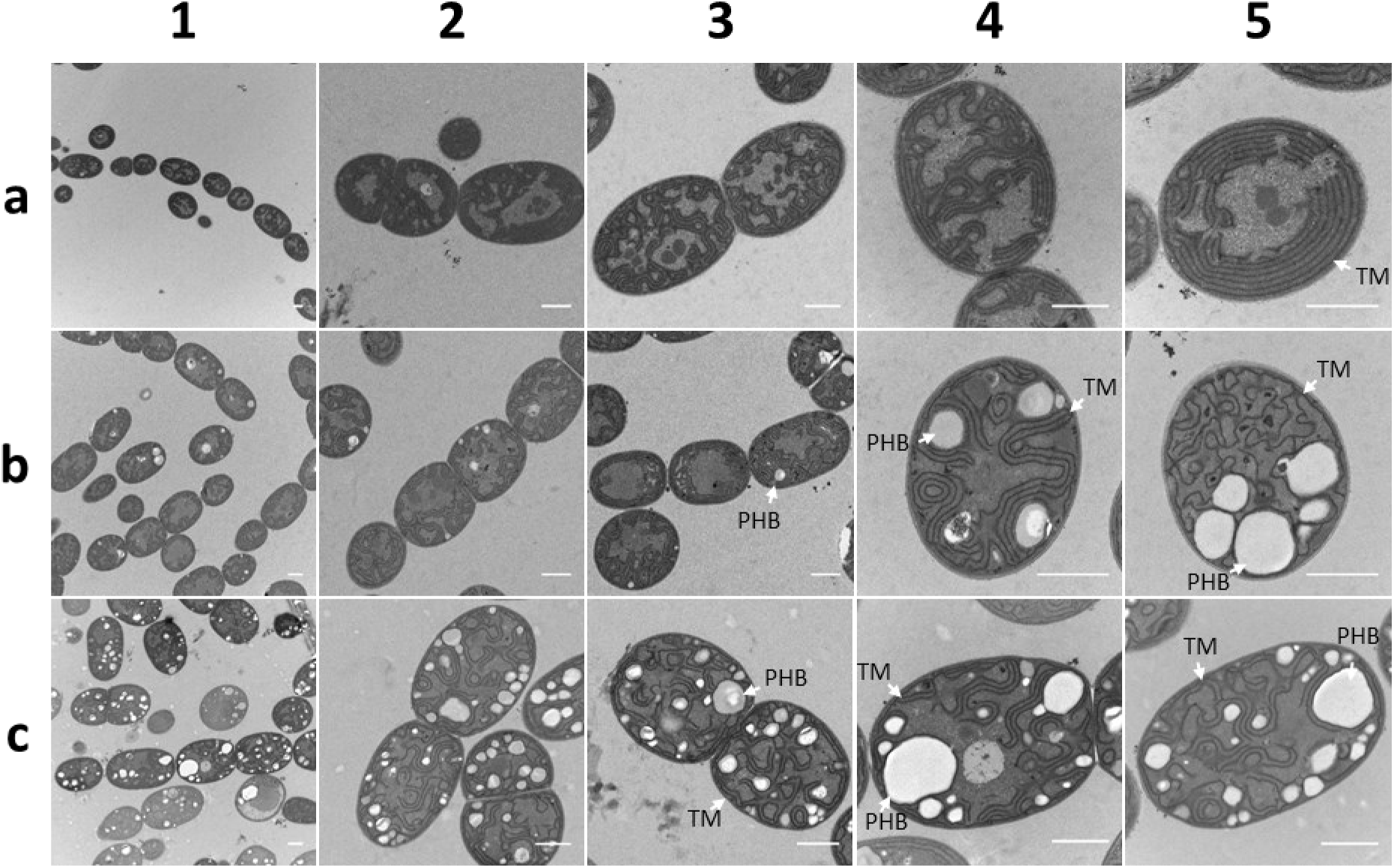
TEM pictures of (**a**) *Nostoc* sp. PCC7120, (**b**) NosPHB2.0, and (**c**) NosPHB3.0 after 12 days of cultivation, TM: Thylakoid membrane, PHB: PHB granule, scale bar = 1 µm.

The TEM pictures confirmed the higher amounts of PHB in NosPHB3.0 than in NosPHB2.0 (Fig. 12 b+c, 1). PHB granule sizes varied for both recombinant strains. In NosPHB2.0, filaments with low number of visible PHB granule (Fig. 12 b, 2+3) and single graveyard cells with higher amount of PHB granule (Fig. 12 b, 4+5) could be distinguished which confirmed the former observation of PHB graveyard forming in NosPHB2.0. In NosPHB3.0, PHB accumulation was clearly visible in the filaments (Fig. 12 c, 1-3). Interestingly, the thylakoid membrane structure was slightly disorganised in both recombinant strains where the parallel, densed thylakoid structure got loosened up (Fig. 12, b+c 4). In the case of the graveyard cells of NosPHB2.0, the organized stacks of thylakoid membranes were further disrupted when more PHB was accumulated (Fig. 12 b, 5). Contrastingly, no forms of similar disturbance of thylakoid membrane were observed in NosPHB3.0 (Fig. 12 c, 4+5). It seemed that the atypical thylakoid membrane shapes of the graveyard cells in NosPHB2.0 could be correlated with the size and amount of PHB granules whereas in NosPHB3.0, phasin PhaP1 utilization could prevent major disorganization.

## Discussion

### Positive effects of PhaP1 on cell viability

In this study, constitutive PHB production introduced into *Nostoc* sp. PCC7120 seemed to exert a negative effect on cell viability and could, therefore, be considered a stress inducer. In fact, a previous proteomic analysis of *E. coli* mutants producing recombinant PHB showed an increased expression of stress related proteins such as GroEL, GroES and DnaK being induced by PHB accumulation (64). In view of this, it was proposed that cytosolic proteins could bind to PHB granules unspecifically and therefore trigger additional stress responses (64). Moreover, PHB biosynthesis competes for the precursor acetyl-CoA with major metabolic pathways like the TCA cycle and the fatty acid synthesis (65), thereby affecting the growth behaviour of PHB producing cells.

Hence, the need to stabilize PHB production in a third generation of PHB-producing *Nostoc* mutants was apparent. Improved cell viability was finally achieved through the utilization of phasin PhaP1 from *Cupriavidus necator* H16, which was integrated into the synthetic PHB operon and subsequently the entire novel operon was integrated into the genome of *Nostoc* sp. PCC7120 resulting in the recombinant NosPHB3.0. This resulted in the stabilization of PHB production which is congruent with the current state of research around phasins. More specifically, previous studies on the introduction of phasin PhaP1 into *E. coli* showed positive effects on growth rates and improved stress response to heat and oxidative stress (66). In a different study, the co-expression of the PHB biosynthetic gene cluster *phaBAC* from *Azotobacter* sp. strain FA8 and *phaP* reduced the response of heat shock proteins in *E. coli*. Additionally, due to phasin expression, unspecific protein binding to PHB granules could be reduced (22). Conversely, in the natural PHB producer *Cupriavidus necator* H16, deletion of *phaP1* lead to a decrease in PHB production rate and a PHA-leaky phenotype (22). Another study found heat shock response proteins GroEL and DnaK attached to PHB granules analogous to recombinant PHB producers without phasin expression (21). Similarly, in the present study, the importance of phasin utilization for PHB stabilization was reiterated and phasin PhaP1 was proven an essential contributor to stable PHB production in recombinant *Nostoc* strain NosPHB3.0.

### Thylakoid membrane disorder due to PHB accumulation

Thylakoid membranes are well-structured, internal membrane systems housing proteins of the light harvesting complex, the respiratory and the photosynthetic electron transport (67–69). The specific morphological arrangement of thylakoid membranes varies between different species of cyanobacteria. In filamentous cyanobacteria, the thylakoid membrane tends to be fascicular, forming short segments of hemispherical loops (70), which were also observed in the herein investigated wild type strain *Nostoc* sp. PCC7120 (Fig. 12, a). Contrastingly, the recorded TEM pictures of recombinant *Nostoc* strains showed filaments containing disorganized thylakoid membranes. Here, an interesting tendency was observed correlating the amount of PHB per filament and the degree of disorder characterizing the respective thylakoid membranes. It is not surprising that increased PHB production fills the intracellular space with the produced biopolymer, thus leading to rearrangements within the cell (64). Therefore, higher PHB/CDW ratios would correlate with a higher percentage of intracellular space being occupied by PHB. In fact, the intracellular space of *Cupriavidus necator* H16 might be occupied by PHB up to an amount of over 70 – 90 % PHB/CDW (15,71). However, an increasingly distorted thylakoid membrane, might affect growth rates and cell viability of the recombinant strain. Thus, more investigation has to be continued.

### Constitutive PHB accumulation in NosPHB3.0

Traditionally, nitrogen limitation is one of the best studied conditions for PHB accumulation in PHB-producing cyanobacteria (59,72,73). For industrial application, cyanobacteria are frequently cultivated in a two-step cultivation process. Firstly, cyanobacterial biomass has to be accumulated in a standard cultivation medium (50). Secondly, the standard medium had to be replaced by a medium with limited nitrogen availability in order to initialize nitrogen starvation and, subsequently, PHB production. This constitutes a complicated cultivation approach, which is not economically favourable, but rather the major disadvantage hindering continuous, industrial cultivations of cyanobacteria (74). The latter therefore encounter difficulties in competing with heterotrophic PHB-producing organisms. In recent studies, through the combination of genetic engineering and optimized cultivation conditions of the cyanobacterial strain *Synechocystis* sp. 6803 PPT1, PHB production rates of up to 60 % *(w/w)* PHB/CDW could be reported. Therein, acetate was described to constitute an organic carbon source able to boost PHB production in cyanobacteria, thus, up to 80 % *(w/w)* PHB/CDW could be achieved (39). However, more sustainable strategies to produce PHB are preferred. Moreover, in the study in question, PHB production was still contingent upon nitrogen and phosphate starvation. Therefore, a system of constitutive PHB production independent of nitrogen-limiting conditions in cyanobacteria would be advantageous regarding industrial upscale cultivations (75). Some cases of cyanobacteria producing PHB under natural phototrophic growth conditions have been described but have not been found preferable for efficient PHB production. More specifically, some studies have reported PHB/CDW rates of only up to 8.5 % *(w/w)* after 21 days of continuous cultivation in the filamentous cyanobacterium *Nostoc muscorum* (72). Likewise, in *Synechocystis* sp. 6803, only small percentage of PHB/ CDW has been reported (43).

In a recent study, phototrophic PHB production was genetically engineered in the non-producer *Synechococcus elongatus* UTEX2973 (76,77). Intriguingly, the utilized PHB operon *phaCAB* also originates from *Cupriavidus necator* H16, similar to the operon used in the present study. However, Roh et al. constructed an inducible PHB production system by setting the operon under the control of the inducible promotor P*_trc_* . Furthermore, no phasins were utilized for the stabilization of PHB biosynthesis in the study in question. Optimized growth conditions (continuous high light of 200 µE m^-2^s^-1^) and the supplementation of 5 % *(v/v)* CO_2_ brought PHB production in the recombinant *Synecococcus elongatus* UTEX 2973 up to 16.7 % PHB/CDW after 10 days of cultivation. Similar PHB production rates were achieved in the present study in cultivations of the first generation of recombinant strains NosPHB1.0, where up to 17 % *(w/w)* PHB/CDW after 8 days of cultivation were achieved. It is not surprising that Roh et al. detected stable PHB production in *Synechococcus elongatus* UTEX2973 due to the inducible regulation system. In constructing an inducible system, the PHB-associated metabolic burden we observed might have been reduced in the recombinant *Synecococcus* strain. However, prolonged cultivation and PHB production in the recombinant *Synecococcus* strain might lead to reduced PHB levels over time, as seen in NosPHB1.0 and NosPHB2.0. Furthermore, the need to supplement cultures with appropriate inducers renders the process more complicated and is thus a disadvantage in comparison with the constitutive system described in the present work. In another recent study, the PHB operon *phaCAB* originated from *Cupriavidus necator* H16 was overexpressed under the constitutively promotor P*_cpc560_* in *Synechococcus elongatus* UTEX2973 (78). Here, up to 7.6 % *(w/w)* PHB/CDW was achieved after 10 days of the same phototropic growth conditions as described before. PHB accumulation in recombinant *Synecococccus* strain was increased to an amount of 17.2 % *(w/w)* PHB/CDW after 10 days via acetate feeding.

In this study, for the first time, successful constitutive PHB production was shown in a recombinant *Nostoc* sp. PCC7120 strain. Although PHB production was reduced in the first mutant generations NosPHB1.0 and NosPHB2.0, the utilization of phasin PhaP1 enabled stable, constitutive PHB production. The herein achieved constitutive PHB production, which was additionally successfully decoupled from nitrogen starvation constitutes an essential step towards sustainable, industrial PHB production.

### Accumulation of filaments lacking PHB and fluorescence signal in recombinant *Nostoc* strains

As previously described, the first two Nostoc mutant generations NosPHB1.0 and NosPHB2.0 were characterized by an abundance of filaments lacking PHB production. Contrastingly, the most recent recombinant *Nostoc* strain NosPHB3.0 displayed a stable PHB production of up to 30% PHB/CDW via the utilization of phasin PhaP1. Nevertheless, even in the latest optimized strain a few empty filaments lacking fluorescence signals corresponding to PHB were still observed. The multifunctional role of PHB precursor acetyl-CoA might prove disadvantageous for overproducer strains of PHB (65). Therefore, it would not be surprising that during cultivation, non-fluorescent filaments could be detected in the culture volume due to precursor bottlenecks.

One of the most intriguing yet underexplored issue of working with cyanobacteria is the occurrence of genetic instability (79). Incidents of genetic instability have been described for ethylene and mannitol overproduction in cyanobacteria. In the former case, the ethylene producing strain was overgrown by a non-producing mutant strain (80). In the latter example, the biosynthesis of mannitol was obstructed by a single frame-shifting mutation leading to the abolishment of mannitol biosynthesis (81). The problem of genetic instability might explain the few filaments found in NosPHB3.0, which were found to lack PHB. For instance, single point mutations in the PHB operon could lead to loss of the ability to produce PHB. Mutations in the promoter sequence P*_psbA_* might lead to reduced transcription rates of translatable mRNA for constitutive protein expression. Single point mutations in the PHB operon and, more specifically, the biosynthetic genes *phaA*, *phaB* or *phaC* could lead to a frameshift resulting in malfunctioning biosynthetic enzymes. Frameshift mutations in phasin phaP1 might negate its stabilizing effect leading to a loss of PHB production similar to NosPHB1.0 and NosPHB2.0.

## Conclusion

This study showed for the first time the recombinant, continuous PHB accumulation in *Nostoc* sp. 7120. PHB accumulation was stabilized by the utilization of phasin PhaP1 from *Cupriavidus necator* H16. Moreover, PhaP1 promoted the formation of PHB granules inside cyanobacterial filaments. Production rates of up to 30 % *(w/w)* PHB/CDW were achieved in the recombinant strain NosPHB3.0.

## Declaration

### Ethics approval and consent to participate

Not applicable

### Consent for publication

Not applicable

### Availability of data and materials

The authors declare that the data supporting the findings of this study are available within the paper and its Supplementary Information files. Should any raw data files be needed in another format they are available from the corresponding author upon reasonable request.

### Competing interests

The authors declare that they have no competing interests

### Funding

This Project was supported by Korea Evaluation Institute of Industrial Technology (KEIT) grant funded by the Korean government (MOTIE) (No. RS-2022-00155902) and Basic Science Research Program through the National Research Foundation of Korea (NRF) funded by the Ministry of Education (2021R1I1A3055799) and by BMBF Grant PHBMax (031B1399B).

### Authors’ contributions

Initial conceptualization PF, JHK and KF; Methodology and interpretation of results, PF and KF; Investigation, PF, CM, Funding acquisition, KF and JHK; Writing original draft preparation, PF and KF; Writing-review & editing, PF and KF; Supervision KF; All authors read and approved the final manuscript.

## Acknowledgements

We thank Andreas Kulik for the support on helping with the initial validation of the PHB quantification method and Christoph Mayer for supporting the HPLC analytics. We thank Iris Maldener for help with *Nostoc* sp. PCC7120 genetics and Peter Wolk for providing pRL-plasmids pRL1049 and pRL271.

## Notes

### Competing Interest Statement

The authors have declared no competing interest.

## References

1. Jambeck JR, Geyer R, Wilcox C, Siegler TR, Perryman M, Andrady A, et al. Plastic waste inputs from land into the ocean. Science. 2015 Feb 13;347(6223):768–71.

2. Shen M, Huang W, Chen M, Song B, Zeng G, Zhang Y. (Micro)plastic crisis: Un-ignorable contribution to global greenhouse gas emissions and climate change. Journal of Cleaner Production. 2020 May 1;254:120138.

3. Bellini S, Tommasi T, Fino D. Poly(3-hydroxybutyrate) biosynthesis by *Cupriavidus necator*: A review on waste substrates utilization for a circular economy approach. Bioresource Technology Reports. 2022 Feb 1;17:100985.

4. Z. Naser A, Deiab I, M. Darras B. Poly(lactic acid) (PLA) and polyhydroxyalkanoates (PHAs), green alternatives to petroleum-based plastics: a review. RSC Advances. 2021;11(28):17151– 96.

5. Ansari S, Fatma T. Cyanobacterial Polyhydroxybutyrate (PHB): Screening, Optimization and Characterization. PLOS ONE. 2016 Jun 30;11(6):e0158168.

6. Koller M, Maršálek L, de Sousa Dias MM, Braunegg G. Producing microbial polyhydroxyalkanoate (PHA) biopolyesters in a sustainable manner. New Biotechnology. 2017 Jul 25;37:24–38.

7. Reinecke F, Steinbüchel A. Ralstonia eutropha Strain H16 as Model Organism for PHA Metabolism and for Biotechnological Production of Technically Interesting Biopolymers. Journal of Molecular Microbiology and Biotechnology. 2008 Oct 29;16(1–2):91–108.

8. Vandamme P, Coenye T. Taxonomy of the genus Cupriavidus: a tale of lost and found. International Journal of Systematic and Evolutionary Microbiology. 2004;54(6):2285–9.

9. Davis DH, Doudoroff M, Stanier RY, Mandel M. Proposal to reject the genus Hydrogenomonas: Taxonomic implications. International Journal of Systematic and Evolutionary Microbiology. 1969;19(4):375–90.

10. Yabuuchi E, Kosako Y, Yano I, Hotta H, Nishiuchi Y. Transfer of Two Burkholderia and An Alcaligenes Species to Ralstonia Gen. Nov.: Proposal of Ralstonia pickettii (Ralston, Palleroni and Doudoroff 1973) Comb. Nov., Ralstonia solanacearum (Smith 1896) Comb. Nov. and Ralstonia eutropha (Davis 1969) Comb. Nov. MICROBIOLOGY and IMMUNOLOGY. 1995;39(11):897–904.

11. Vaneechoutte M, Kämpfer P, De Baere T, Falsen E, Verschraegen G. Wautersia gen. nov., a novel genus accommodating the phylogenetic lineage including Ralstonia eutropha and related species, and proposal of Ralstonia [Pseudomonas] syzygii (Roberts et al. 1990) comb. nov. International Journal of Systematic and Evolutionary Microbiology. 2004;54(2):317–27.

12. Obruca S, Marova I, Snajdar O, Mravcova L, Svoboda Z. Production of poly(3-hydroxybutyrate- co-3-hydroxyvalerate) by Cupriavidus necator from waste rapeseed oil using propanol as a precursor of 3-hydroxyvalerate. Biotechnol Lett. 2010 Dec 1;32(12):1925–32.

13. Schlegel HG, Gottschalk G. Poly-β-hydroxybuttersäure, ihre Verbreitung, Funktion und Biosynthese. Angewandte Chemie. 1962;74(10):342–7.

14. Holmes PA. Applications of PHB - a microbially produced biodegradable thermoplastic. Physics in Technology. 1985 Jan;16(1):32–6.

15. Schlegel HG, Gottschalk G, Von Bartha R. Formation and Utilization of Poly-β-Hydroxybutyric Acid by Knallgas Bacteria (Hydrogenomonas). Nature. 1961 Jul;191(4787):463–5.

16. Peoples OP, Sinskey AJ. Poly-β-hydroxybutyrate biosynthesis in Alcaligenes eutrophus H16: Characterization of the genes encoding β-ketothiolase and acetoacetyl-CoA reductase *. Journal of Biological Chemistry. 1989 Sep 15;264(26):15293–7.

17. Peoples OP, Sinskey AJ. Poly-β-hydroxybutyrate (PHB) biosynthesis in Alcaligenes eutrophus H16: Identification and characterization of the PHB polymerase gene (phbC) *. Journal of Biological Chemistry. 1989 Sep 15;264(26):15298–303.

18. Pohlmann A, Fricke WF, Reinecke F, Kusian B, Liesegang H, Cramm R, et al. Genome sequence of the bioplastic-producing “Knallgas” bacterium Ralstonia eutropha H16. Nat Biotechnol. 2006 Oct;24(10):1257–62.

19. Pfeiffer D, Jendrossek D. Localization of Poly(3-Hydroxybutyrate) (PHB) Granule-Associated Proteins during PHB Granule Formation and Identification of Two New Phasins, PhaP6 and PhaP7, in Ralstonia eutropha H16. Journal of Bacteriology. 2012 Oct 8;194(21):5909–21.

20. Jendrossek D. Polyhydroxyalkanoate Granules Are Complex Subcellular Organelles (Carbonosomes). Journal of Bacteriology. 2009 May;191(10):3195.

21. Pötter M, Müller H, Steinbüchel A. Influence of homologous phasins (PhaP) on PHA accumulation and regulation of their expression by the transcriptional repressor PhaR in Ralstonia eutropha H16. Microbiology. 2005;151(3):825–33.

22. Wieczorek R, Pries A, Steinbüchel A, Mayer F. Analysis of a 24-kilodalton protein associated with the polyhydroxyalkanoic acid granules in Alcaligenes eutrophus. Journal of Bacteriology. 1995 May;177(9):2425–35.

23. York GM, Stubbe J, Sinskey AJ. The Ralstonia eutropha PhaR Protein Couples Synthesis of the PhaP Phasin to the Presence of Polyhydroxybutyrate in Cells and Promotes Polyhydroxybutyrate Production. J Bacteriol. 2002 Jan;184(1):59–66.

24. Bresan S, Jendrossek D. New Insights into PhaM-PhaC-Mediated Localization of Polyhydroxybutyrate Granules in Ralstonia eutropha H16. Appl Environ Microbiol. 2017 May 31;83(12):e00505–17.

25. Pfeiffer D, Jendrossek D. PhaM Is the Physiological Activator of Poly(3-Hydroxybutyrate) (PHB) Synthase (PhaC1) in Ralstonia eutropha. Appl Environ Microbiol. 2014 Jan;80(2):555– 63.

26. Jendrossek D, Handrick R. Microbial Degradation of Polyhydroxyalkanoates. Annu Rev Microbiol. 2002 Oct;56(1):403–32.

27. Chandani N, Mazunder PB, Bhattacharjee A. Production of Polyhydroxybutyrate (biopolymer) by Bacillus tequilensis NCS-3 Isolated from Municipal Waste Areas of Silchar, Assam. 2012;3(12).

28. Nikel PI, Pettinari MJ, Galvagno MA, Méndez BS. Poly(3-Hydroxybutyrate) Synthesis by Recombinant Escherichia coli arcA Mutants in Microaerobiosis. Appl Environ Microbiol. 2006 Apr;72(4):2614–20.

29. Van Wegen RJ, Ling Y, Middelberg APJ. Industrial Production of Polyhydroxyalkanoates Using *Escherichia Coll*: An Economic Analysis. Chemical Engineering Research and Design. 1998 Mar 1;76(3):417–26.

30. Choi J, Lee SY. Economic considerations in the production of poly(3-hydroxybutyrate-co-3- hydroxyvalerate) by bacterial fermentation. Appl Microbiol Biotechnol. 2000 Jun 1;53(6):646–9.

31. Ryu HW, Hahn SK, Chang YK, Chang HN. Production of poly(3-hydroxybutyrate) by high cell density fed-batch culture of Alcaligenes eutrophus with phospate limitation. Biotechnology and Bioengineering. 1997;55(1):28–32.

32. Kulpreecha S, Boonruangthavorn A, Meksiriporn B, Thongchul N. Inexpensive fed-batch cultivation for high poly(3-hydroxybutyrate) production by a new isolate of *Bacillus megaterium*. Journal of Bioscience and Bioengineering. 2009 Mar 1;107(3):240–5.

33. Lopez-Arenas T, González-Contreras M, Anaya-Reza O, Sales-Cruz M. Analysis of the fermentation strategy and its impact on the economics of the production process of PHB (polyhydroxybutyrate). Computers & Chemical Engineering. 2017 Dec 5;107:140–50.

34. Sirohi R, Prakash Pandey J, Kumar Gaur V, Gnansounou E, Sindhu R. Critical overview of biomass feedstocks as sustainable substrates for the production of polyhydroxybutyrate (PHB). Bioresource Technology. 2020 Sep 1;311:123536.

35. Costa JAV, Moreira JB, Lucas BF, Braga V da S, Cassuriaga APA, Morais MG de. Recent Advances and Future Perspectives of PHB Production by Cyanobacteria. Industrial Biotechnology. 2018 Oct;14(5):249–56.

36. Koch M, Forchhammer K. Polyhydroxybutyrate: A Useful Product of Chlorotic Cyanobacteria. Microbial Physiology. 2021 May 12;31(2):67–77.

37. Singh AK, Mallick N. Advances in cyanobacterial polyhydroxyalkanoates production. FEMS Microbiology Letters. 2017 Nov 1;364(20):fnx189.

38. Troschl C, Meixner K, Drosg B. Cyanobacterial PHA Production—Review of Recent Advances and a Summary of Three Years’ Working Experience Running a Pilot Plant. Bioengineering. 2017 Jun;4(2):26.

39. Koch M, Bruckmoser J, Scholl J, Hauf W, Rieger B, Forchhammer K. Maximizing PHB content in Synechocystis sp. PCC 6803: a new metabolic engineering strategy based on the regulator PirC. Microbial Cell Factories. 2020 Dec 22;19(1):231.

40. Oliver NJ, Rabinovitch-Deere CA, Carroll AL, Nozzi NE, Case AE, Atsumi S. Cyanobacterial metabolic engineering for biofuel and chemical production. Current Opinion in Chemical Biology. 2016 Dec 1;35:43–50.

41. Gupta V, Ratha SK, Sood A, Chaudhary V, Prasanna R. New insights into the biodiversity and applications of cyanobacteria (blue-green algae)—Prospects and challenges. Algal Research. 2013 Mar 1;2(2):79–97.

42. Carr NG. The occurrence of poly-β-hydroxybutyrate in the blue-green alga, Chlorogloea fritschii. Biochimica et Biophysica Acta (BBA) - Biophysics including Photosynthesis. 1966 Jun 8;120(2):308–10.

43. Kaewbai-ngam A, Incharoensakdi A, Monshupanee T. Increased accumulation of polyhydroxybutyrate in divergent cyanobacteria under nutrient-deprived photoautotrophy: An efficient conversion of solar energy and carbon dioxide to polyhydroxybutyrate by *Calothrix scytonemicola* TISTR 8095. Bioresource Technology. 2016 Jul 1;212:342–7.

44. Panda B, Mallick N. Enhanced poly-β-hydroxybutyrate accumulation in a unicellular cyanobacterium, Synechocystis sp. PCC 6803. Letters in Applied Microbiology. 2007 Feb 1;44(2):194–8.

45. Panda B, Jain P, Sharma L, Mallick N. Optimization of cultural and nutritional conditions for accumulation of poly-β-hydroxybutyrate in *Synechocystis* sp. PCC 6803. Bioresource Technology. 2006 Jul 1;97(11):1296–301.

46. Sharma L, Mallick N. Accumulation of poly-β-hydroxybutyrate in *Nostoc muscorum*: regulation by pH, light–dark cycles, N and P status and carbon sources. Bioresource Technology. 2005 Jul 1;96(11):1304–10.

47. Monshupanee T, Nimdach P, Incharoensakdi A. Two-stage (photoautotrophy and heterotrophy) cultivation enables efficient production of bioplastic poly-3-hydroxybutyrate in auto-sedimenting cyanobacterium. Sci Rep. 2016 Nov 15;6(1):37121.

48. Carpine R, Olivieri G, Hellingwerf KJ, Pollio A, Marzocchella A. Industrial Production of Poly- β-hydroxybutyrate from CO2: Can Cyanobacteria Meet this Challenge? Processes. 2020 Mar;8(3):323.

49. Lachance MAndré. Genetic Relatedness of Heterocystous Cyanobacteria by Deoxyribonucleic Acid-Deoxyribonucleic Acid Reassociation. International Journal of Systematic and Evolutionary Microbiology. 1981;31(2):139–47.

50. Rippka R, Deruelles J, Waterbury JB, Herdman M, Stanier RY. Generic Assignments, Strain Histories and Properties of Pure Cultures of Cyanobacteria. Microbiology. 1979;111(1):1–61.

51. Tamas I, Svircev Z, Andersson SG. Determinative value of a portion of the nifH sequence for the genera Nostoc and Anabaena (cyanobacteria). Curr Microbiol. 2000 Sep;41(3):197–200.

52. Elhai J, Vepritskiy A, Muro-Pastor AM, Flores E, Wolk CP. Reduction of conjugal transfer efficiency by three restriction activities of Anabaena sp. strain PCC 7120. Journal of Bacteriology. 1997 Mar;179(6):1998–2005.

53. Kaneko T, Nakamura Y, Wolk CP, Kuritz T, Sasamoto S, Watanabe A, et al. Complete genomic sequence of the filamentous nitrogen-fixing cyanobacterium Anabaena sp. strain PCC 7120. DNA Res. 2001 Oct 31;8(5):205–13; 227–53.

54. Menestreau M, Rachedi R, Risoul V, Foglino M, Latifi A. The CcdB toxin is an efficient selective marker for CRISPR-plasmids developed for genome editing in cyanobacteria. MicroPubl Biol. 2022;2022:10.17912/micropub.biology.000512.

55. Wolk CP, Vonshak A, Kehoe P, Elhai J. Construction of shuttle vectors capable of conjugative transfer from Escherichia coli to nitrogen-fixing filamentous cyanobacteria. Proceedings of the National Academy of Sciences. 1984 Mar;81(5):1561–5.

56. Chen M, Li J, Zhang L, Chang S, Liu C, Wang J, et al. Auto-flotation of heterocyst enables the efficient production of renewable energy in cyanobacteria. Sci Rep. 2014 Feb 6;4(1):3998.

57. Bertani G. Lysogeny at Mid-Twentieth Century: P1, P2, and Other Experimental Systems. J Bacteriol. 2004 Feb;186(3):595–600.

58. Gibson DG, Young L, Chuang RY, Venter JC, Hutchison CA, Smith HO. Enzymatic assembly of DNA molecules up to several hundred kilobases. Nat Methods. 2009 May;6(5):343–5.

59. Schlebusch M, Forchhammer K. Requirement of the Nitrogen Starvation-Induced Protein Sll0783 for Polyhydroxybutyrate Accumulation in Synechocystis sp. Strain PCC 6803. Applied and Environmental Microbiology. 2010 Sep 15;76(18):6101–7.

60. Taroncher-Oldenburg G, Nishina K, Stephanopoulos G. Identification and analysis of the polyhydroxyalkanoate-specific beta-ketothiolase and acetoacetyl coenzyme A reductase genes in the cyanobacterium Synechocystis sp. strain PCC6803. Appl Environ Microbiol. 2000 Oct;66(10):4440–8.

61. Elhai J. Strong and regulated promoters in the cyanobacterium Anabaena PCC 7120. FEMS Microbiology Letters. 1993 Dec 1;114(2):179–84.

62. Olmedo-Verd E, Muro-Pastor AM, Flores E, Herrero A. Localized Induction of the ntcA Regulatory Gene in Developing Heterocysts of Anabaena sp. Strain PCC 7120. Journal of Bacteriology. 2006 Sep 15;188(18):6694–9.

63. Wunschiers R, Axelsson R, Lindblad P. Effects of Growth on Dinitrogen on the Transcriptome and Predicted Proteome of Nostoc PCC 7120 [Internet]. arXiv; 2006 [cited 2024 Aug 23]. Available from: http://arxiv.org/abs/q-bio/0604031

64. Han MJ, Yoon SS, Lee SY. Proteome Analysis of Metabolically EngineeredEscherichia coli Producing Poly(3-Hydroxybutyrate). Journal of Bacteriology. 2001 Jan;183(1):301–8.

65. Lee SY, Chang HN. Production of poly(3-hydroxybutyric acid) by recombinant Escherichia coli strains: genetic and fermentation studies. Can J Microbiol. 1995 Dec 15;41(13):207–15.

66. de Almeida A, Catone MV, Rhodius VA, Gross CA, Pettinari MJ. Unexpected Stress-Reducing Effect of PhaP, a Poly(3-Hydroxybutyrate) Granule-Associated Protein, in Escherichia coli▿. Appl Environ Microbiol. 2011 Sep;77(18):6622–9.

67. Berger S, Ellersiek U, Steinmüller K. Cyanobacteria contain a mitochrondrial complex I- homologous NADH-dehydrogenase. FEBS Letters. 1991 Jul 29;286(1):129–32.

68. Stürzl E, Scherer S, Böger P. Interaction of respiratory and photosynthetic electron transport, and evidence for membrane-bound pyridine-nucleotide dehydrogenases in *Anabaena variabilis*. Physiologia Plantarum. 1984 Apr;60(4):479–83.

69. Mullineaux CW. Co-existence of photosynthetic and respiratory activities in cyanobacterial thylakoid membranes. Biochimica et Biophysica Acta (BBA) - Bioenergetics. 2014 Apr;1837(4):503–11.

70. Mareš J, Strunecký O, Bučinská L, Wiedermannová J. Evolutionary Patterns of Thylakoid Architecture in Cyanobacteria. Front Microbiol [Internet]. 2019 Feb 22 [cited 2024 Oct 1];10. Available from: https://www.frontiersin.org/journals/microbiology/articles/10.3389/fmicb.2019.00277/full

71. Mravec F, Obruca S, Krzyzanek V, Sedlacek P, Hrubanova K, Samek O, et al. Accumulation of PHA granules in Cupriavidus necator as seen by confocal fluorescence microscopy. FEMS Microbiology Letters. 2016 May 1;363(10):fnw094.

72. Bhati R, Mallick N. Production and characterization of poly(3-hydroxybutyrate-co-3- hydroxyvalerate) co-polymer by a N2-fixing cyanobacterium, Nostoc muscorum Agardh. Journal of Chemical Technology & Biotechnology. 2012;87(4):505–12.

73. de Philippis R, Sili C, Vincenzini M. Glycogen and poly-β-hydroxybutyrate synthesis in Spirulina maxima. Microbiology. 1992;138(8):1623–8.

74. Drosg B, Fritz I, F G, Silvestrini L. Photo-autotrophic Production of Poly(hydroxyalkanoates) in Cyanobacteria. Chemical and Biochemical Engineering Quarterly. 2015 May 27;29.

75. Price S, Kuzhiumparambil U, Pernice M, Ralph PJ. Cyanobacterial polyhydroxybutyrate for sustainable bioplastic production: Critical review and perspectives. Journal of Environmental Chemical Engineering. 2020 Aug 1;8(4):104007.

76. Roh H, Lee JS, Choi HI, Sung YJ, Choi SY, Woo HM, et al. Improved CO2-derived polyhydroxybutyrate (PHB) production by engineering fast-growing cyanobacterium *Synechococcus elongatus* UTEX 2973 for potential utilization of flue gas. Bioresource Technology. 2021 May 1;327:124789.

77. Yu J, Liberton M, Cliften PF, Head RD, Jacobs JM, Smith RD, et al. Synechococcus elongatus UTEX 2973, a fast growing cyanobacterial chassis for biosynthesis using light and CO2. Sci Rep. 2015 Jan 30;5(1):8132.

78. Lee SY, Lee JS, Sim SJ. Cost-effective production of bioplastic polyhydroxybutyrate via introducing heterogeneous constitutive promoter and elevating acetyl-Coenzyme A pool of rapidly growing cyanobacteria. Bioresource Technology. 2024 Feb 1;394:130297.

79. Jones PR. Genetic Instability in Cyanobacteria – An Elephant in the Room? Front Bioeng Biotechnol. 2014 May 2;2:12.

80. Takahama K, Matsuoka M, Nagahama K, Ogawa T. Construction and analysis of a recombinant cyanobacterium expressing a chromosomally inserted gene for an ethylene-forming enzyme at the *psbAI* locus. Journal of Bioscience and Bioengineering. 2003 Jan 1;95(3):302–5.

81. Jacobsen JH, Frigaard NU. Engineering of photosynthetic mannitol biosynthesis from CO2 in a cyanobacterium. Metabolic Engineering. 2014 Jan 1;21:60–70.

