## Supplemental Material for "A novel recombinant PHB production platform in filamentous cyanobacteria avoiding nitrogen starvation while preserving cell viability"

**Supplementary information:**

Table S 1: Strains used and created in this study

| Name | Genotype | Reference |
| --- | --- | --- |
| *E. coli* NEB10β | *Δ(ara-leu)7697 araD139 fhuA ΔlacX74 galK16 galE15* $e14^-$ *ϕ80dlacZΔM15 recA1 relA1 endA1 nupG rpsL* ($Sm^R$) *rph spoT1 Δ(mrr-hsdRMS-mcrBC)*, $Sm^R$ | NEB |
| *E. coli* Stellar cells | F- *endA1 supE44 thi-1 recA1 relA1 gyrA96 phoA Φ80dlacZΔM15* Δ(lacZYA-argF) U169 Δ(*mrr-hsdRMS-mcrBC*), Δ*mcrA*, λ- | TAKARA |
| *E. coli* J53/RP4 | RP4, $Ap^R$, $Km^R$, $Tc^R$ | Datta 1971 |

| | | |
|---|---|---|
| *E. coli* HB101 | F– *thi*-1 *hsd*S20($r_B^-$, $m_B^-$) *sup*E44 *rec*A13 *ara*-14 *leu*B6 *pro*A2 *lac*Y1 *gal*K2 *rps*L20 ($Sm^R$) *xyl*-5 *mtl*-1, $Sm^R$ | Promega |
| *E. coli* HB101/ pRL528 | HB101+ pRL528, $Cm^R$, $Sm^R$ | Elhai 1988 |
| *E. coli* HB101/ pRL528/pRL1049-$P_{psbA}$-*phaCAB* | HB101+ pRL528 + pRL1049-$P_{psbA}$-*phaCAB,* $Cm^R$, $Sm^R/Sp^R$ | This study |
| *E. coli* HB101/ pRL528/pPF08 | HB101 + pPF08, $Cm^R$, $Sm^R/Sp^R$, $Em^R$ | This study |
| *E. coli* HB101/ pRL528/pPF10 | HB101 + pPF10, $Cm^R$, $Sm^R/Sp^R$, $Em^R$ | This study |
| *Nostoc* sp. PCC7120 | Wild-type strain | Pasteur culture collection PCC |
| NosPHB1.0 | *Nostoc* sp. PCC7120 with replicative plasmid pRL1049-$P_{psbA}$-*phaCAB*; $Sm^R/Sp^R$ | This study |
| NosPHB2.0 | *Nostoc* sp. PCC7120 with integrated PHB operon ($P_{psbA}$-*phaCAB*) inside neutral site (*nucA*-*nuiA* region), $Sm^R/Sp^R$ | This study |
| NosPHB3.0 | *Nostoc* sp. PCC7120 with integrated PHB operon ($P_{psbA}$-*phaCAB*-*apcBA*(*phaP*)) inside neutral site (*nucA*-*nuiA* region), $Sm^R/Sp^R$ | This study |
| Abbreviations: Ap: Ampicillin, Cm: Chloramphenicol, Sm: Streptomycin, Sp: Spectinomycin, Km: Kanamycin, Tc: Tetracycline | | |

Table S 2: Primer used in this study

| Name | Primer | Description |
|---|---|---|
| pPF08_fragment 1 fw | acattgcagttgagaacccagaagctgctgGCAATGGCAACAACGTTGCGCAAACTATTA | Gibson primer to create DNA fragment 1 for pPF08 |
| pPF08_fragment 1 rev | cagatccttcccacaaaaaaacctcaaatgGATCTAGATATCGAATTTCTGCCATTCATCCG | Gibson primer to create DNA fragment 1 for pPF08 |
| pPF08_fragment 2 fw | GATGAATGGCAGAAATTCGATATCTAGATCcatttgaggtttttttgtgggaaggatctg | Gibson primer to create DNA fragment 2 for pPF08 |
| pPF08_fragment 2 rev | GTCAACCAATATTCATTGAGATCCTCTAGAacacctgataattacctgatggtcaaaaat | Gibson primer to create DNA fragment 2 for pPF08 |
| pPF08_fragment 3 fw | atttttgaccatcaggtaattatcaggtgtTCTAGAGGATCTCAATGAATATTGGTTGACAC | Gibson primer to create DNA fragment 3 for pPF08 |
| pPF08_fragment 3 rev | ctgggaaatcccagtggtgcaacgccaacaGAGTTTGTAGAAACGCAAAAAGGCCATCCG | Gibson primer to create DNA fragment 3 for pPF08 |
| pPF08_fragment 4 fw | CGGATGGCCTTTTTGCGTTTCTACAAACTCtgttggcgttgcaccactgggatttcccag | Gibson primer to create DNA fragment 4 for pPF08 |
| pPF08_fragment 4 rev | TAATAGTTTGCGCAACGTTGTTGCCATTGCcagcagcttctgggttctcaactgcaatgt | Gibson primer to create DNA fragment 4 for pPF08 |
| pPF10_fragment 1 fw | GCCACGGCAAAGAAGACGACGGCTGCCTGAGAGCTCTTGACCGAACGCAGCGGTGGTAAC | Gibson primer to create DNA fragment 1 for pPF10 |
| pPF10_fragment 1 rev | ttccagtttcagcagggaaacagctgcaccTCAGCCCATATGCAGGCCGCCGTTGAGCGA | Gibson primer to create DNA fragment 1 for pPF10 |

Table S 3: Plasmids used in this study

| Name | Characteristics | Reference or Source |
|---|---|---|
| RP4 | R+, met, pro, Tra+ IncP; $Ap^R$, $Km^R$, $Tc^R$ | Datta 1971 |
| pRL528 | Helper plasmid for mobilization used in triparental mating: $Mob_{ColK}$, M.AvaI, M.Eco47II, $Cm^R$ | Elhai et al. 1988 |

| | | |
|---|---|---|
| pRL271 | Vector for triparental conjugation and subsequent *sacB*-mediated positive selection of double crossover mutants in *Nostoc* sp. PCC7120, $Cm^R$, $Em^R$ | Cai et al. 1990 |
| pRL1049 | Self-replicating plasmid, $Sm^R/Sp^R$ | Black et al. 1994 |
| pRL1049-$P_{psbA}$-*phaCAB* | pRL1049 harboring PHB operon $P_{psbA}$-*phaCAB*, $Sm^R/Sp^R$ | Kindly provided by Jörg Scholl |
| pPF08 | pRL271 with homologous regions of *nucA-nuiA* and expanded PHB operon $P_{psbA}$-*phaCAB* with spectinomycin/streptomycin resistance cassette *aad1* and t1t2 terminator, $Sm^R/Sp^R$, $Cm^R$, $Em^R$ | This study |
| pPF10 | pRL271 with homologous regions of *nucA-nuiA* and expanded PHB operon $P_{psbA}$-*phaCAB* with insertion of intergenic region of *abcBA* and *phaP* and spectinomycin/streptomycin resistance cassette *aad1* and t1t2 terminator, $Sm^R/Sp^R$, $Cm^R$, $Em^R$ | This study |
| Abbreviations: Ap: Ampicillin, Cm: Chloramphenicol, Em: Erythromycin, Sm: Streptomycin, Sp: Spectinomycin | | |

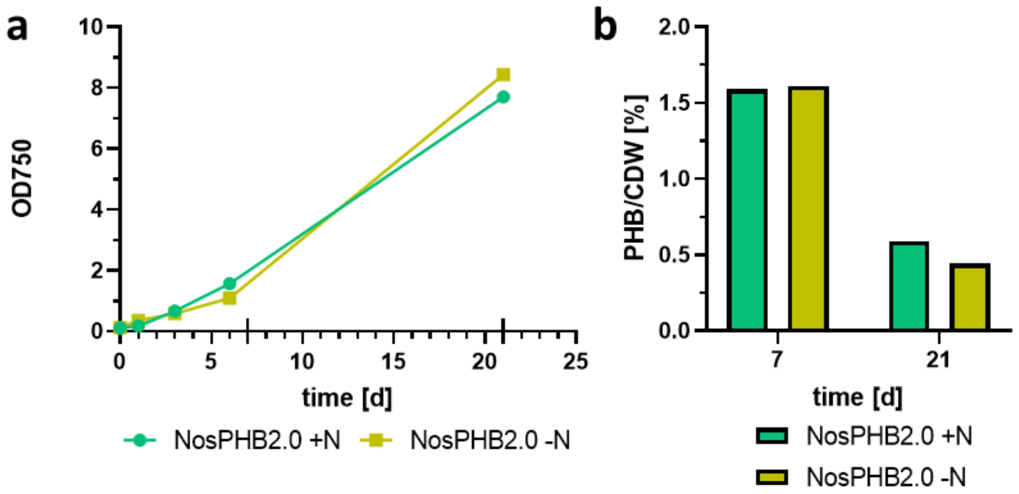


Fig. S 1: Growth experiment and PHB content of NosPHB2.0 **(a)** Growth curve of NosPHB2.0 in BG11 (+N, green) and $BG11_0$ (-N, yellow), Growth was recorded by measuring of the strain's $OD_{750}$, longer tick on x-axis indicates sample taken for PHB quantification. Each point represents one biological experiment **(b)** PHB content of NosPHB2.0 after 7 and 21 days of continuous growth condition in BG11 (+N, green) and $BG11_0$ (-N, yellow). Each data set represents one biological sample.

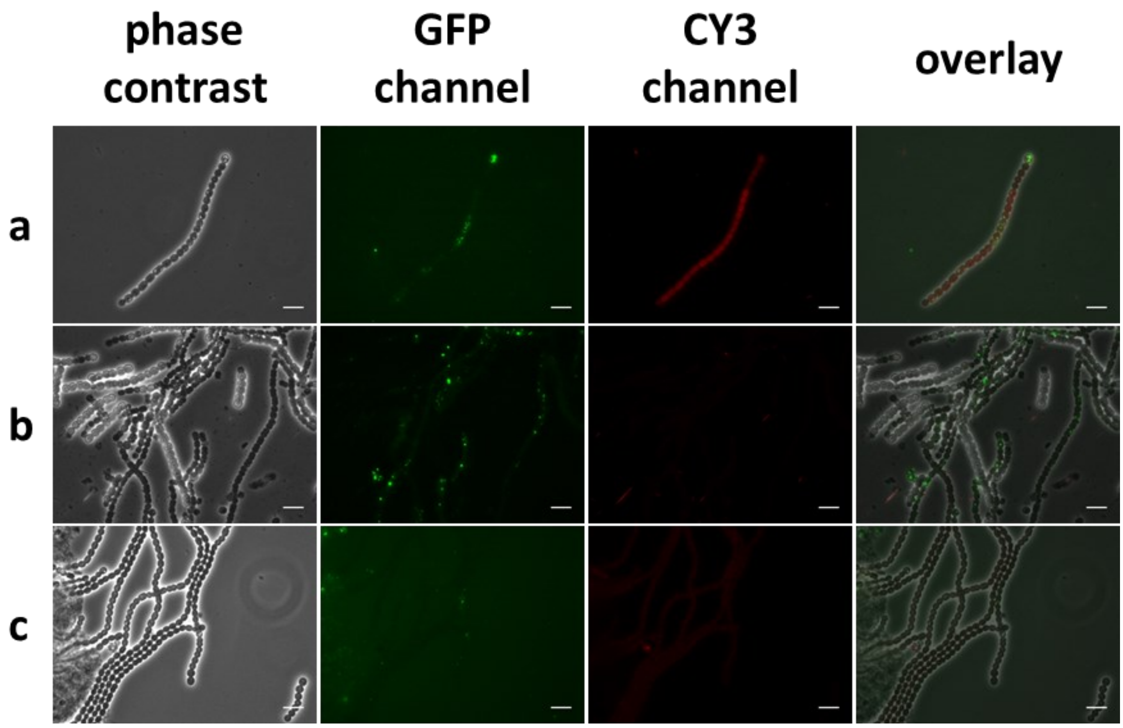


Fig. S 2: Microscopic images of NosPHB2.0 after 3 days of continuous growth condition **(a)** NS: 5 μmol $m^{-2}s^{-1}$, 0 rpm; **(b)** S: 50-60 μmol $m^{-2}s^{-1}$, 120 rpm; **(c)** DS: 20 μmol $m^{-2}s^{-1}$, 120 rpm, PHB granules are visualized by BODIPY staining and detected using the GFP channel PHB granules are visualized with BODIPY staining (GFP channel), autofluorescence (CY3 channel), overlay of phase contrast, GFP and CY3 channel, heterocyst formation was indicated with white arrows,scale bar = 10 μm.

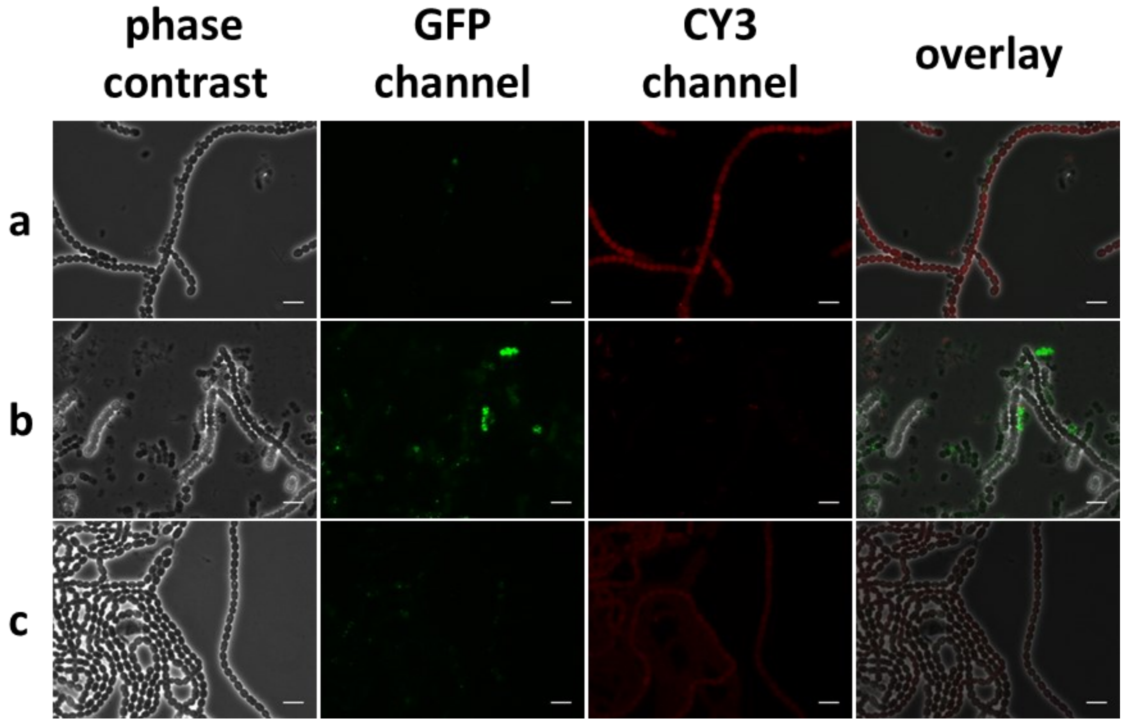


Fig. S 3: Microscopic images of NosPHB2.0 after 7 days of continuous growth condition **(a)** NS: 5 µmol $m^{-2}s^{-1}$, 0 rpm; **(b)** S: 50-60 µmol $m^{-2}s^{-1}$, 120 rpm; **(c)** DS: 20 µmol $m^{-2}s^{-1}$, 120 rpm, PHB granules are visualized by BODIPY staining and detected using the GFP channel PHB granules are visualized with BODIPY staining (GFP channel), autofluorescence (CY3 channel), overlay of phase contrast, GFP and CY3 channel, heterocyst formation was indicated with white arrows, scale bar = 10 µm.

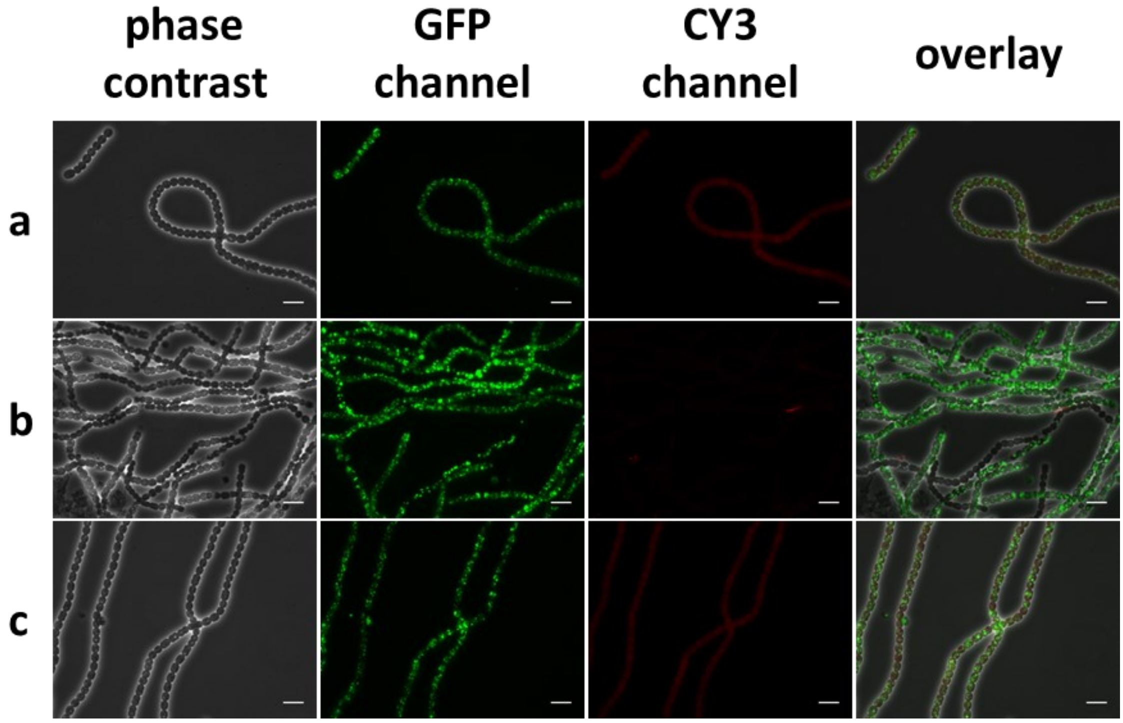


Fig. S 4: Microscopic images of NosPHB3.0 after 3 days of continuous growth condition **(a)** NS: 5 μmol $m^{-2}s^{-1}$, 0 rpm; **(b)** S: 50-60 μmol $m^{-2}s^{-1}$, 120 rpm; **(c)** DS: 20 μmol $m^{-2}s^{-1}$, 120 rpm, PHB granules are visualized by BODIPY staining and detected using the GFP channel PHB granules are visualized with BODIPY staining (GFP channel), autofluorescence (CY3 channel), overlay of phase contrast, GFP and CY3 channel, heterocyst formation was indicated with white arrows, scale bar = 10 μm.

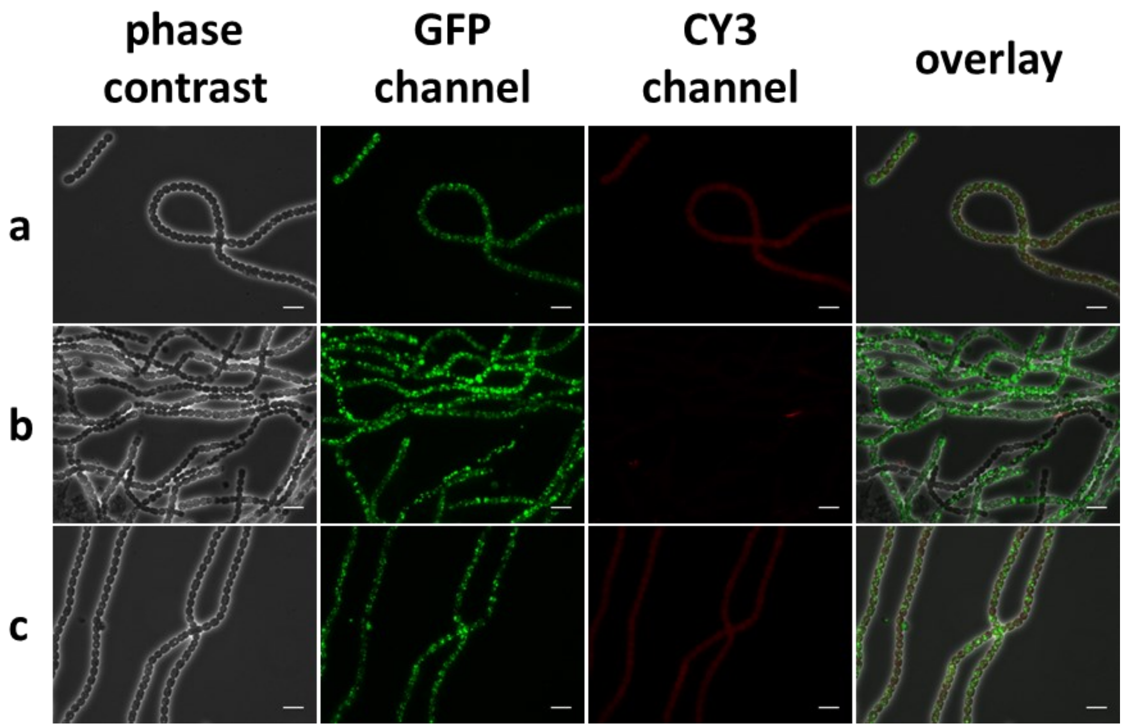


Fig. S 5: Microscopic images of NosPHB3.0 after 7 days of continuous growth condition **(a)** NS: 5 µmol $m^{-2}s^{-1}$, 0 rpm; **(b)** S: 50-60 µmol $m^{-2}s^{-1}$, 120 rpm; **(c)** DS: 20 µmol $m^{-2}s^{-1}$, 120 rpm, PHB granules are visualized by BODIPY staining and detected using the GFP channel PHB granules are visualized with BODIPY staining (GFP channel), autofluorescence (CY3 channel), overlay of phase contrast, GFP and CY3 channel, heterocyst formation was indicated with white arrows, scale bar = 10 µm.
